# Optogenetic interrogation of the lateral-line sensory system reveals mechanisms of pattern separation in the zebrafish brain

**DOI:** 10.1101/2025.02.07.637118

**Authors:** Nicolas Velez-Angel, Sihao Lu, Brian Fabella, Caleb C. Reagor, Holland R. Brown, Yuriria Vázquez, Adrian Jacobo, A. J. Hudspeth

## Abstract

The ability of animals to interact with their environment hinges on the brain’s capacity to distinguish between patterns of sensory information and accurately attribute them to specific sensory organs. The mechanisms by which neuronal circuits discriminate and encode the source of sensory signals remain elusive. To address this, we utilized as a model the posterior lateral line system of larval zebrafish, which is used to detect water currents. This system comprises a series of mechanosensory organs called neuromasts, which are innervated by neurons from the posterior lateral line ganglion. By combining single-neuromast optogenetic stimulation with whole-brain calcium imaging, we developed a novel approach to investigate how inputs from neuromasts are processed. Upon stimulating individual neuromasts, we observed that neurons in the brain of the zebrafish show diverse selectivity properties despite a lack of topographic organization in second-order circuits. We further demonstrated that complex combinations of neuromast stimulation are represented by sparse ensembles of neurons within the medial octavolateralis nucleus (MON) and found that neuromast input can be integrated nonlinearly. Our approach offers an innovative method for spatiotemporally interrogating the zebrafish lateral line system and presents a valuable model for studying whole-brain sensory encoding.

## Introduction

The ability to discriminate complex patterned stimuli is crucial for animals to respond effectively to a changing environment. To differentiate sensory cues, central sensory systems must therefore adopt diverse computational strategies tailored to their anatomical structures^1,2^. To achieve this, sensory cortices typically either map sensory receptor space topographically^3^ or integrate receptor inputs into pseudo-random ensembles^4–6^ that rely on the emergent properties of population activity^7–9^.

Topographic mapping enhances the efficiency of sensory coding but is typically constrained to larger brains that are able to fit the space of input parameters. This constraint is evident in structures such as ocular dominance columns in the primary visual cortex of some mammals^10^, which encode a narrow range of features^11^. As the combinatorial space of stimuli becomes more complex, such as in the encoding of natural scenes^12,13^, neuronal ensembles employ other strategies to efficiently discriminate between inputs. These strategies include coding through nonlinear summation of inputs, decorrelation across patterned stimuli, and sparse neuronal ensembles^2,7,14^.

The lateral line system of fish and amphibians is a mechanoreceptive system that detects water currents^15^ and enables those organisms to navigate their environment, avoid obstacles, escape predators, detect prey, and conduct behaviors such as rheotaxis and schooling^16–19^. Because this sensory system must accurately differentiate across a broad array of stimuli, a critical question arises: how do central circuits in the zebrafish brain process inputs from the posterior lateral line (pLL) to discriminate various flow patterns?

Neuromasts, the sensory receptors in the pLL, are specialized epithelial organs with mechanoreceptive hair cells^20^ that detect the origin, frequency, velocity, and direction of water currents. Each neuromast typically contains 10-20 hair cells^21^ arranged in opposing orientations along the anteroposterior (AP) or dorsoventral (DV) axis^22,20^. These neuromasts are arranged in a grid-like pattern across the body’s AP axis, which may provide spatial information about the location and origin of nearby cues_23,24_.

Our understanding of how discrete neuromast inputs are organized within the central nervous systems of larval zebrafish is limited. Afferent neurons in the pLL ganglion (pLLg) project to the medial octavolateralis nucleus (MON) in the hindbrain^25^. Single-unit recordings from goldfish suggest that many cells in the MON are insensitive to flow direction^24^ and display receptive fields of varying sizes, some of which span widely separated neuromasts^23,24^. Furthermore, single-cell labeling shows that, in addition to the MON, afferent neurons from the pLL innervate neurons in the eminentia granularis of the cerebellum^26^ as well as Mauthner cells.^27,28^ It is unclear whether these projections are present in zebrafish larvae. As denoted by the zebrafish atlas, the MON also sends projections to the torus semicircularis (TS) and optic tectum (OT) of the midbrain^29–31^ and to the superior medulla.^30^ Single-unit responses in these areas, particularly the TS, show heterogeneous temporal patterns and selectivity to the location of objects moving along the AP axis^32^. Little is known about how a hydrodynamic stimulus’ position, distance, and speed are encoded in the zebrafish’s brain^33^.

The simple circuitry and stereotyped neuromast arrangement of the larval pLL make it an ideal model for exploring the integration of sensory stimuli and the associated computational mechanisms. Although this approach is promising, a major challenge in studying the pLL is the difficulty of precisely stimulating individual neuromasts and recording the resulting neuronal activity. Flow-based stimulation techniques lack the fine control to target single neuromasts. They often generate turbulence upon interacting with the neuromast’s hydrodynamic boundary layer and cannot activate multiple neuromasts simultaneously.

Using the optical accessibility of the zebrafish larva, we have developed a system to interrogate individual neuromasts in the pLL by optogenetic stimulation. Evidence shows that the opening of light-gated ion channels in hair cells can induce in pLL ganglion afferents spiking activity comparable to that elicited by mechanical stimulation^28^. Applying optogenetics to single neuromasts can therefore provide the precision and flexibility to discreetly activate combinations of neuromasts in any imaginable order, a strategy that overcomes the limitations of hydrodynamic flow-based stimuli. Combining optogenetic stimulation with calcium imaging through single-objective light-sheet microscopy^34,35^, we created a scalable system to deliver varying spatiotemporal patterns of neuromast input and simultaneously assess their effects on brain-wide neuronal activity. Using this novel approach, we ask how neuromast position is encoded in the zebrafish’s brain and establish this sensory organ as a model for research on sensory computation.

## Results

### Single-neuromast optogenetics

Neuromasts in the pLL of a larval zebrafish are linearly arranged along the AP axis, spaced at least 100 µm apart (Figure 1A). Positioned symmetrically on both sides of the body, they are ideally suited for precise and individualized optogenetic stimulation. We developed a laser-scanning system with a galvanometric optical relay to target neuromasts in zebrafish larvae expressing CoChR-GFP (Figure 1B, Figures S1 and S2). With this system, we targeted specific labeled neuromasts along the tail of a zebrafish larva six days after fertilization (6 dpf), either individually or in various combinations (Figure 1C), with negligible spillover to neighboring targets (Figure S4). Because the transition between irradiation of different locations within the fish required less than 500 μs, faster than the activation kinetics of the channelrhodopsin^36^, we could activate multiple neuromasts almost simultaneously.

**Figure 1:**
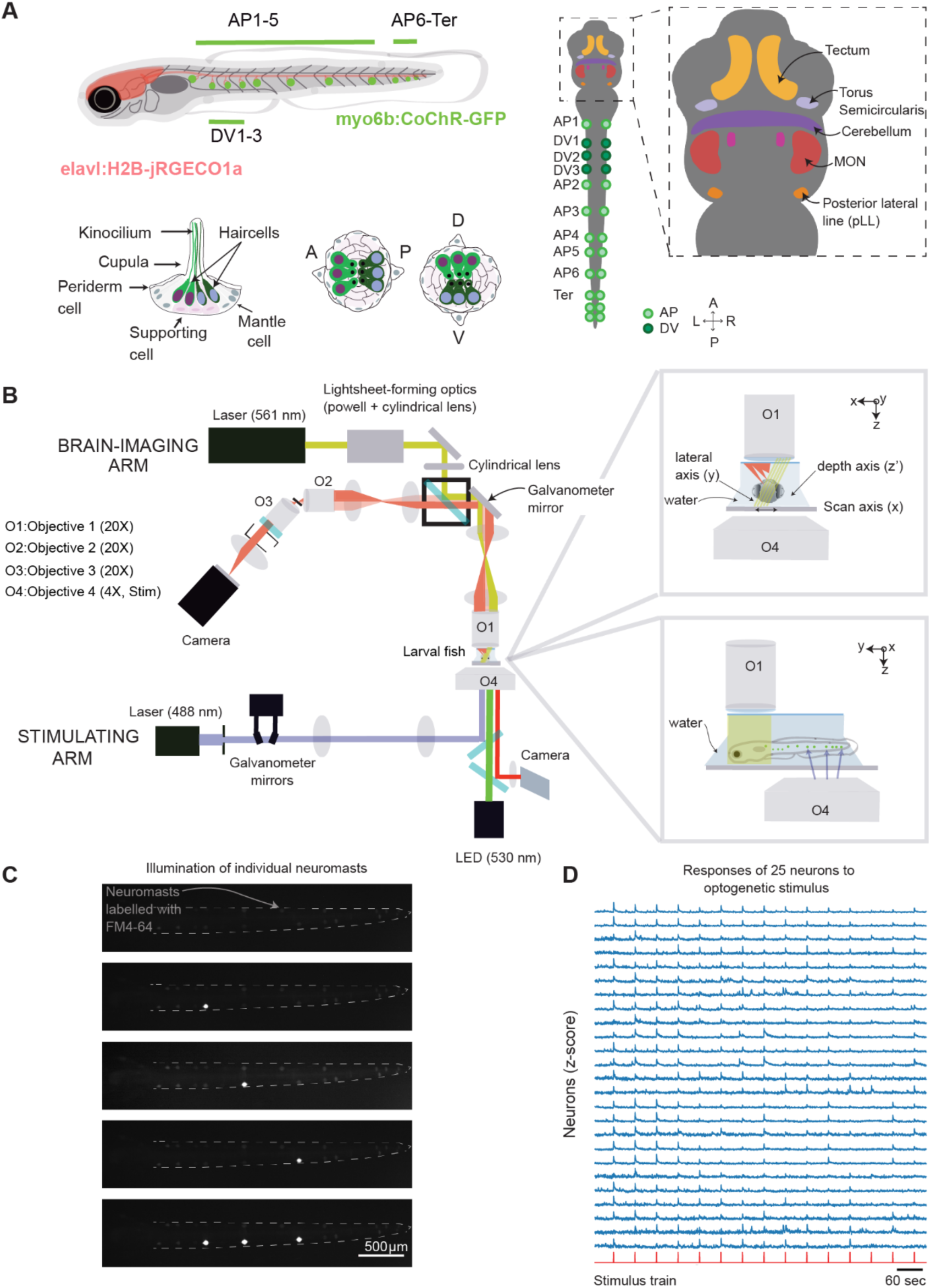
Dual optogenetic stimulation of single neuromasts and whole-brain imaging in the larval zebrafish. **A**. On the left, a schematic diagram illustrates the posterior lateral line (pLL) of a 6-day-old larval zebrafish, which has approximately 11-12 neuromasts (green circles). Eight or nine of these neuromasts exhibit hair cells polarized along the anterior-posterior (AP) axis (AP1-AP6), with two or three terminal neuromasts at the tail. In addition, three neuromasts—typically located between AP1 and AP2—have hair cells polarized along the dorsal-ventral (DV) axis. All neuromasts connect to the brain through axons that extend from the pLL ganglion. On the right, a schematic diagram depicts the regions of the brain that have previously been associated with responses to pLL stimuli. Projections from the pLL innervate the MON, which then connects to the other areas. **B.** A schematic diagram depicts the dual optical system used to optogenetically activate individual neuromasts during whole-brain imaging in larval zebrafish. The upper part shows the geometry of the SCAPE microscope used to perform whole-brain imaging from above. The bottom panel shows the laser-scanning system positioned at the larva’s tail to target labeled neuromasts in the pLL for optogenetic stimulation. The top inset on the right shows that, as the light sheet of the SCAPE microscope scans across the lateral dimension of the larva, it can capture fluorescence along an axis nearly perpendicular to the light sheet. The lower inset depicts the lateral view of the apparatus. The objective lenses of the SCAPE and optogenetic system are offset laterally by 1 mm, which allows them to target the head and tail of the larva, respectively, without interference. **C.** The top panel provides an example image of a 6 dpf larva’s tail with neuromasts labeled with FM4-64. In the second to fourth panels, the bright spot are caused by the illumination of the laser which precisely targeted individual neuromasts along the larval trunk. In the bottom panel, all three neuromasts were activated almost simultaneously. **D**. The z–scored calcium-imaging traces of 25 neurons responsive to optogenetic stimulation of one neuromast show that our system can activate single neuromasts and observe neurons that activate in response to this stimulus. Neuronal calcium signals are shown in blue; the timings of the optogenetic stimuli are represented in orange at the bottom of the traces.

To integrate optogenetic stimulation with whole-brain calcium imaging, we used a SCAPE microscope^34^ to image the larva’s brain (Figure S3). The optogenetic optical pathway was positioned approximately 1 mm behind the SCAPE’s objective, to target the larva’s tail while the SCAPE imaged the brain. In *Tg(myo6b:CoChR-GFP); Tg(elavl1:H2B-jRGECO1a)* fish, we found that neurons in the brain responded reliably to the optogenetic stimulation of a neuromast at powers exceeding 26.2 mW·mm^-2^, above which sensitivity plateaued (Figure S2, n = 4-6). We observed neurons that responded to one neuromast with high fidelity across trials (Figure 1D). Our system thus efficiently stimulates individual neuromasts in *Tg(myo6b:CoChR-GFP); Tg(elav1:H2B-jRGECO1a*) larvae and integrates seamlessly with whole-brain calcium imaging.

### Organization of neurons to stimulation of individual neuromasts

By systematically stimulating individual neuromasts along one side of a larva with our optogenetic system, we explored how individual neuromasts are represented in the brain (Figure 2A). Most neurons that responded to individual neuromast input were located at the ipsilateral MON (Figure 2B). This response was highly lateralized, for we observed very few neurons in the contralateral MON despite documented presence of excitatory and inhibitory projections from one MON to its contralateral counterpart^30^. Responsive neurons were also found in the superior contralateral medulla, likely associated with the anterior rhombencephalic turning region (ARTR)^37^ or the mesencephalic locomotor region (MLR)^38^ and other areas in the inferior hindbrain. In addition, few neurons were observed in the ipsilateral cerebellum, the contralateral TS, and the OT (Figure S5), which are regions previously associated with encoding hydrodynamic flow.

**Figure 2:**
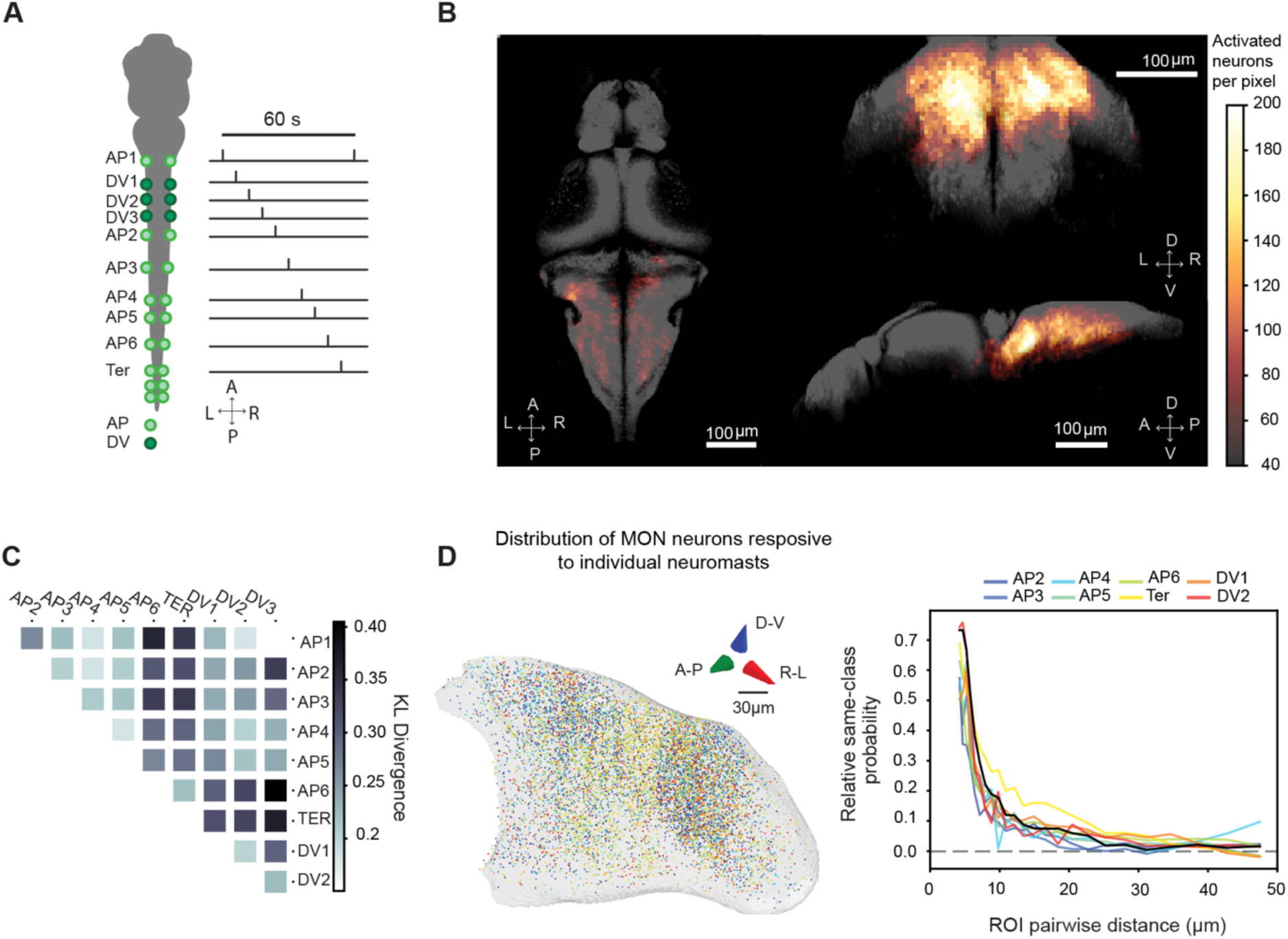
Optogenetic stimulation of single neuromasts. **A.** In an illustration of the stimulation protocol for individual neuromasts. Six to ten neuromasts were stimulated along the left side of each fish. Each neuromast was stimulated every 60 s. **B.** The locations of neurons responsive to optogenetic neuromast stimulation in 25 fish co-registered to the MapZeBrain atlas. Images show the ipsilateral MON and superior medulla as major anatomical sites for neuronal responsiveness. They also show distributed activities across the hindbrain. Left panel: Horizontal maximal projection. A: Anterior; P, Posterior; L, Left; R, Right. Right upper panel: Coronal maximal projection. D, Dorsal; V, Ventral; L, left; R, right. Lower right panel: Sagittal maximal projection. D, Dorsal; V, Ventral, A: Anterior; P: Posterior. **C.** A matrix depicting the Kullback-Liebler divergence (KL divergence) average of the spatial distributions of neurons responsive to each of the neuromasts of the pLL. The matrix relates the distribution of spatial neurons responding to different neuromasts and shows that neurons responding to terminal neuromasts differ in their overall location from other neurons responding to other neuromasts of the pLL. **D.** On the left, a 3-D rendering of the MON displays the spatial distribution of neurons responding to optogenetic activation from the 8 most prominent neuromasts across samples. The scale bar is 30 μm. On the right, cluster homogeneity analysis is used to study the probability of finding two neurons responsive to the same neuromast at increasing distances from each other within the MON. The graph shows that the probability of encountering a neuron responsive to a neuromast of the same identity is highest at distances less than 10 μm.

Many second-order sensory areas organize sensory input through topographic maps^3^. We sought to determine whether a topographic map exists in the MON. After filtering for single selective neurons in the MON, we observed no clear topographic organization between neuromasts along the AP axis and no apparent difference in the distribution of neurons responding to AP *versus* DV neuromasts (Figure S6).

Even if a global spatial organization of tuning is not apparent^39^, neuronal organization might still occur at a more local scale. To analyze this, we compared the distributions of neurons responsive to each neuromast within the MON. We found that the anatomical distribution of terminal neuromasts, including AP6, differs from that of trunk neuromasts. Both AP and DV neuromasts showed a more homogeneous distribution between them (Figure 2C). Terminal (Ter) neuromasts seem to be scattered more broadly across the MON, whereas trunk neuromasts are concentrated in two specific areas with visible subclusters of neurons activated by the same stimulus (Figure 2D)^40^. Moreover, when using a nearest-neighbor algorithm to assess the cluster homogeneity between neurons responsive to the same neuromast (Figure 2D), we found fields of neurons that responded to the same trunk neuromasts at separations as great as about 10-20 µm.

Although we find that there is a global distinction between neuromasts in the trunk (both AP and DV) and those near the tail, neurons responsive to stimulation of individual neuromasts do not seem to be topographically organized within the MON.

### Selectivity of neurons to individual neuromast stimulation

The lack of a fine-grained topographic map does not hinder the ability of a sensory system to discriminate among patterned stimuli^11^. In many cases, if neurons are selective to either one or a specific group of inputs, sensory stimuli can nonetheless be encoded faithfully. Therefore, the selectivity patterns of neurons might suffice to encode neuromast location and identity. When stimulating different sets of neuromasts while performing whole-brain imaging (Figure 3A), we observed both single-selective and mixed-selective neurons responding to individual neuromast input (Figure 3B, Figure S7).

**Figure 3:**
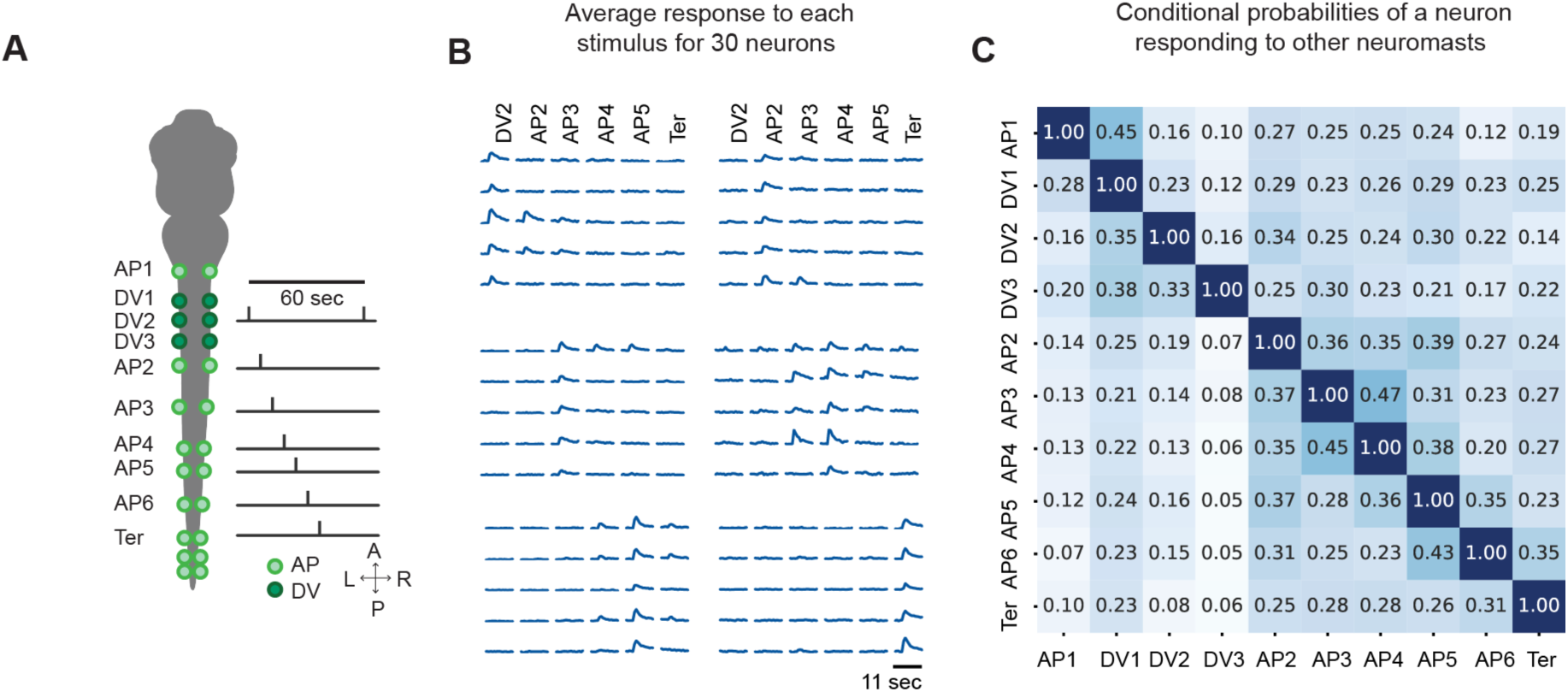
Neurons activated by individual neuromast stimulation show both single and mixed selectivity properties. **A**. A schematic diagram depicts our neuromast stimulation protocol in which 5 to 9 individual neuromasts were stimulated along the left side of the fish. **B.** The z-scored mean activity of neurons is shown for one larva that responded to the optogenetic stimulation of each neuromast shown in A. Traces serve as an example to convey that neurons are both single and mixed-selective to the activation of an individual neuromast and that selectivity follows the body axis of the fish. **C**. A matrix that shows the conditional probability of when neurons responding to one neuromast are also active when stimulating other neuromasts. For example, row 1 of the heatmap depicts the likelihood of a neuron activated by AP1 stimulation but also responds to stimulation of DV1, DV2, AP2, AP3, and so on.

To determine whether there is spatial organization within neuronal selectivity, we calculated the conditional probability of a neuron’s responding to a given neuromast while also responding to any other neuromast (Figure 3C). Through this approach, we found that many responsive neurons are single-selective. However, we also observed the presence of mixed-selective neurons. Neurons responding to multiple neuromasts were more likely to integrate inputs from neighboring neuromasts, indicating a non-random pattern of mixed selectivity that followed the body axis of the zebrafish. As the distance between neuromasts increased, the probability of a neuron’s firing in response to both neuromasts decreased.

These results suggest strategic encoding of the AP axis in the larva’s body through the patterns of neuromast selectivity. The data support the notion that the fish’s brain contains neurons with receptive fields of variable sizes^29^.

### Linear decodability of the neuronal population to individual neuromast stimulation

We wondered whether the neuronal activity from the stimulation of individual neuromasts can discriminate neuromast identity on a trial-by-trial basis. Using a support vector machine (SVM) with a linear kernel, we classified individual neuromast input with up to 90% accuracy based on the binarized responses of neurons correlating to neuromast input (Figure 4A-C).

**Figure 4.**
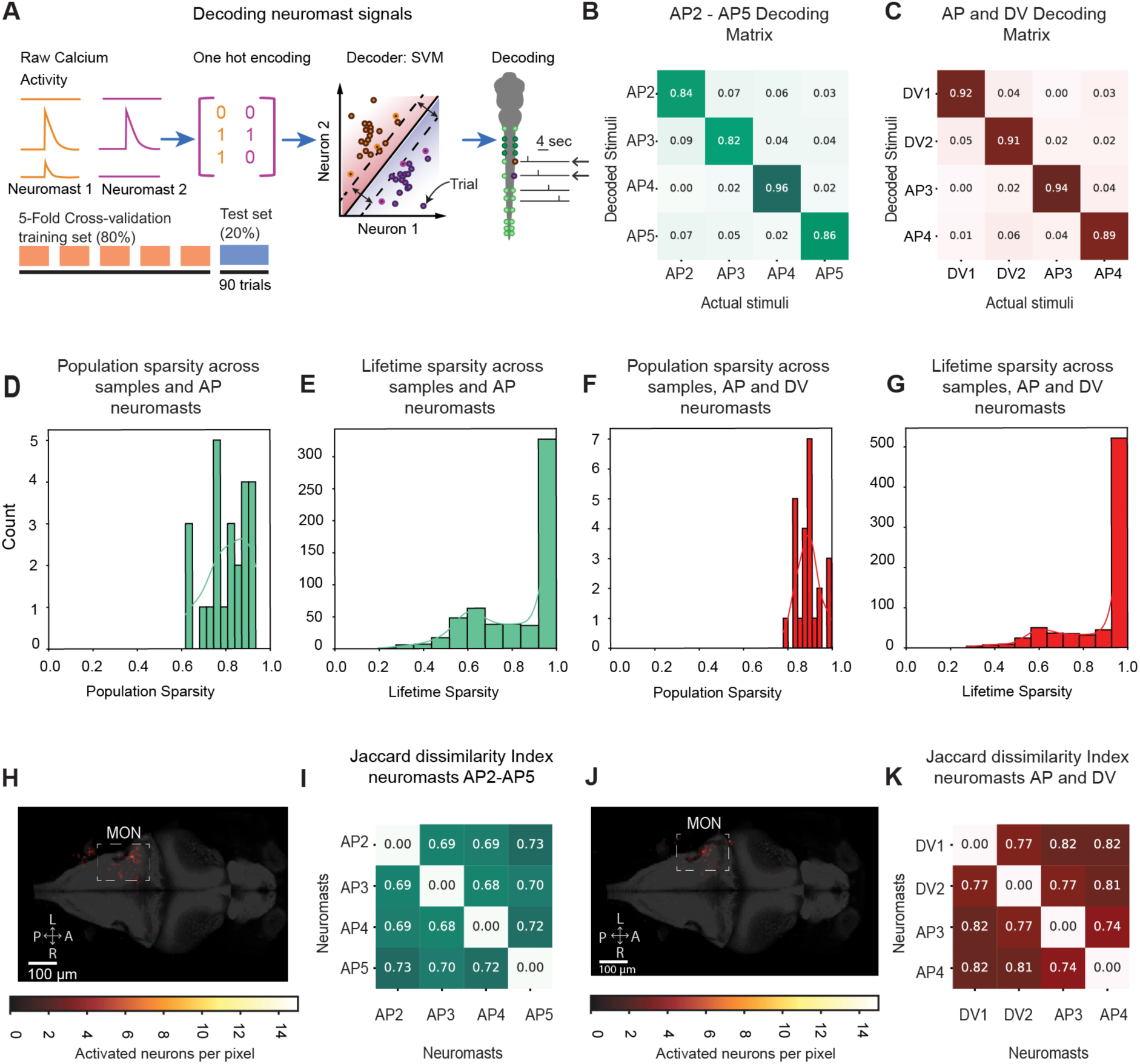
Neuromast identity can be decoded by a linear classifier acting on the population of neuromast-responsive neurons. **A**. A diagram depicts our strategy to decode neuromast input from neuronal activity. Using a peri-stimulus window of 2 s for each trial to train a linear support vector machine (SVM), we binarize calcium transients occurring during optogenetic stimulation. The consequent model is used to predict the origin of the neuromast input. **B**. The average confusion matrix displays the decoding accuracy of the neuromast input between AP neuromasts (AP2-AP5, n=6 fish). **C**. The average confusion matrix displays the classification accuracy of neuronal activity elicited when stimulating AP and DV neuromasts (DV1, DV2, AP3, AP4, n=6 fish). **D.** A histogram shows the distribution of sparseness values of the rows of the weight matrix from the SVM classifier in B. We denominate this metric as the population sparsity because it relates to the amount of neuromast-responsive neurons that participate in the decoding. A value of 1 suggests that one neuronal weight carries all the information and 0 shows equal neuronal weights. **E.** The distribution of sparseness values of the non-zero weights across the columns for the SVM classifier in B. relates to how many neuromast categories each neuronal weight helps decode. A value of 1 means a neuronal weight helps decode only one stimulus class, whereas lower values suggest equal contribution. **F.** The same analysis as in D. is shown but for the SVM classifier in C. **G.** The same as in E. is shown but for the SVM classifier in C. **H.** The location of neurons with weights greater than three standard deviations above the mean across classes shows most decoding neurons are in MON. **I**. The Jaccard dissimilarity index of the weights across each neuromast class for the SVM classifier in E. shows that non-overlapping ensembles of neurons are involved in decoding each neuromast class. **J**. The same as in K. is shown but for the weights in F. **K**. The same as in L. is shown but for weights in F.

By examining the decoder’s weight distribution, we explored the mechanisms of pattern separation for neuromast identity. Theoretical work suggests that neural networks can better distinguish overlapping signals when neurons represent inputs sparsely^6,41,42^–across neuromast categories—population sparseness—and across neurons—lifetime sparseness. Population sparseness indicates how sparse information is represented across the population of neurons, with a value of 1 suggesting that one neuron carries all the information and 0 indicating equal participation from all neurons. For both decoders, population sparseness values tended towards 1, a result that suggests that neuromast input was encoded by a small ensemble of neurons (Figure 4D and 4F). The mean population sparseness for neuromasts AP2-AP5 was 0.8 ± 0.04 (mean, SEM, n=6), whereas for neuromasts DV1, DV2, AP3, and AP4, it was 0.89 ± 0.02 (mean ± SEM, n = 6).

Lifetime sparseness, on the other hand, reveals mixed selectivity: a value of 1 means a neuron decodes only one stimulus class, whereas lower values suggest equal contribution.

After removing neurons that showed all weights at zero, we observed that most neurons had a lifetime sparsity of 1. This result suggests that single selectivity (Figure 4E and 4G) and weights were differently distributed across categories (Figure 4H and 4J). Weights significantly different from zero were predominantly in the lateral MON, with a few neurons also in ipsilateral medial regions near cerebellar and medullary structures (Figure 4I and 4K).

When decoding neuromast input on a selected population of only mixed selective neurons, we find that these neurons are still able to decode, albeit to a lower extent, the identity of the neuromast being stimulated (Figure S8). Moreover, this decrease in accuracy is accompanied by a drop in the population and lifetime sparseness measurements of the decoder weights. These results show that neuromast identity can be decoded by small ensembles of mainly single-selective neurons, with only a small contribution from mixed-selective neurons to provide robustness to the system.

### Linear decodability of the neuronal population to combinatorial neuromast stimulation

Although individual sensory inputs from neuromasts can be decoded by linear classifiers, it remains unclear whether more complex combinations of neuromasts can be decoded. To investigate this, we first activated all possible pairs of neuromasts between AP2 and AP5. We found responsive neurons in the MON, both in anterior and posterior parts, as well as the contralateral superior medulla, the TS, and OT (Figure S9).

Using principal-component analysis (PCA) to explore the geometry of the responsive population we then analyzed the mean activity of responsive neurons. Differences in paired stimuli were most apparent in the second, third, and fourth components, which explained 8%, 6%, and 5% of the variance respectively. Across these three principal components, we observed distinct manifolds for all six pairs of stimulated neuromasts (Figure 5B and Figure S10 for individual examples). Trajectories showed maximum separation immediately after the optogenetic stimulus. Trajectories from pairs with non-overlapping sets of neuromasts were oriented in opposite directions, whereas those with one common neuromast appeared to be nearly orthogonal in this low-dimensional space.

**Figure 5:**
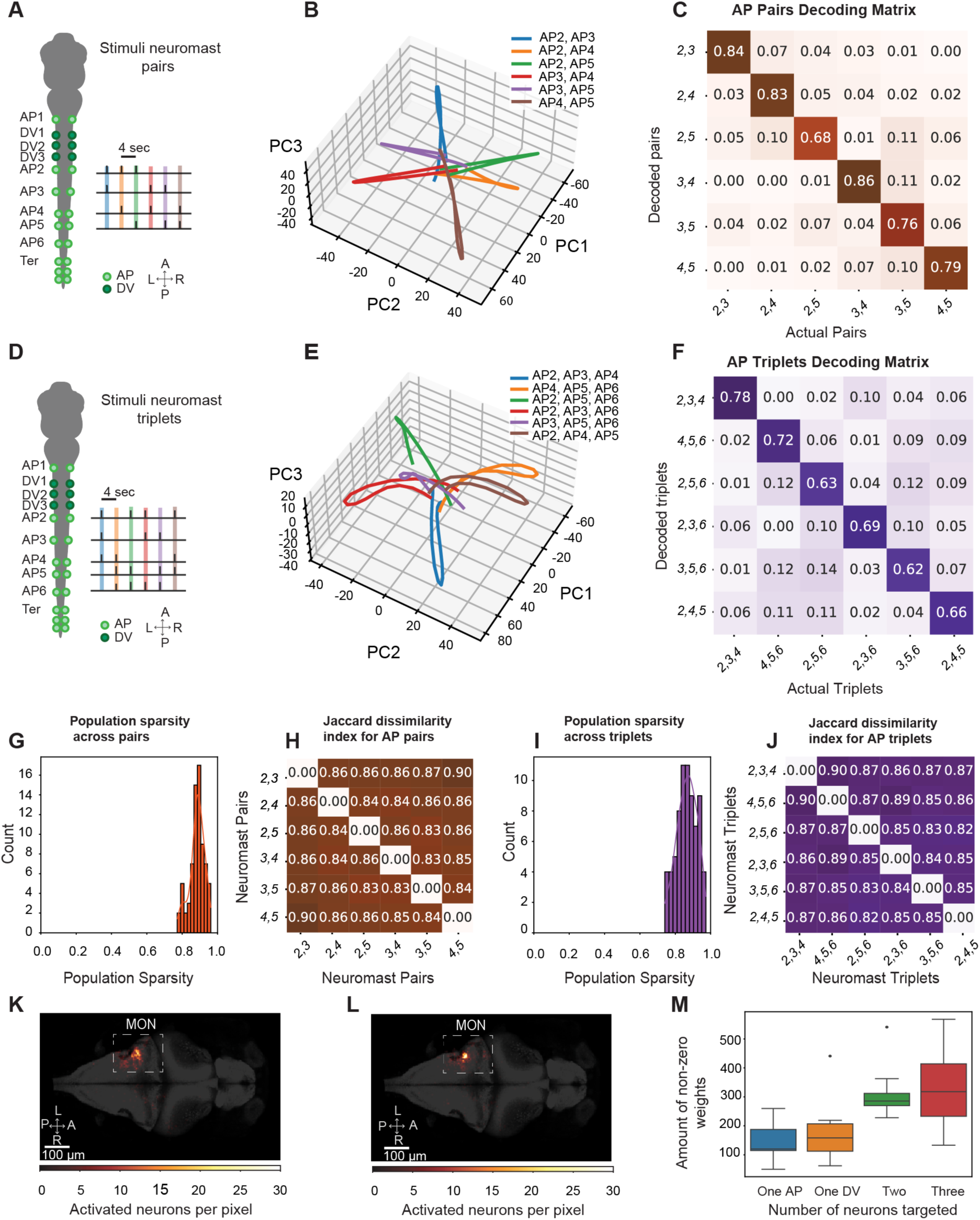
Combinations of neuromast stimulation patterns are represented by population activity (n=12 fish). **A.** In an illustration of the stimulation protocol for pairs of neuromasts, pairs of neuromasts from AP2 to AP5 are stimulated with an inter-stimulus interval of 4 s. **B.** The mean PCA projection of the trial-averaged neural activity in the fish onto to the second to fourth PC components reveals distinct trajectories for each stimulated neuromast pair. **C.** The average confusion matrix displays the decoding accuracy of paired neuromast inputs across fish samples. **D.** In an illustration of the stimulation protocol for selected neuromast triples between AP2 and AP6, six combinations were selected corresponding to AP2-AP3-AP4, AP4-AP5-AP6, AP2-AP3-AP6, AP2-AP5-AP6, AP2-AP4-AP5, AP3-AP4-AP6. **E.** PCA projection of the trial-averaged neural activity onto the second to fourth PCs reveals separate trajectories for each stimulated neuromast triplet. **F.** The average confusion matrix for tripled stimuli displays the average decoding accuracy of tripled stimuli across fish samples. **G.** The distribution of sparseness values from the weights in C) across stimulus classes is depicted, indicating high levels of sparsity in the neuromast-responsive population. **H.** The Jaccard dissimilarity index of weights across neuromast class for the SVM classifier in C) relates encoding of each neuromast identity by different ensembles of neurons. **I.** The distribution of sparseness values from the weights in F) across stimulus classes is depicted. **J.** The Jaccard dissimilarity index of the weights across neuromast class for the SVM classifier in F) is shown. **K-L**. The location of neurons with weights greater than three standard deviations above the mean across classes shows most decoding neurons are in the MON. **M.** A graph compares the number of non-zero weights of each SVM across the number of neuromasts of the stimulus paradigms for single (Figure 4), double and triple stimuli (Figure 5).

Divergent trajectories after dimensionality reduction suggest that pattern separation is achievable for similar neuromast combinations. To confirm this, we applied a linear decoder as used in Figure 4. The decoder achieved high discriminability across neuromast pairs, with average accuracies at 0.8 ± 0.03 (mean ± SEM, n = 12, Figure 5C). Accuracy values were slightly lower compared to single neuromast stimulation and showed relative uniformity across classes.

We extended our experimental analysis to neuromast triplets by selecting six triplets from neuromasts AP2 to AP6, ensuring that AP6 was separated from the terminal neuromasts. These triplets were chosen to effectively cover the combinatorial stimulus space of the stimulated neuromasts (Figure 5D). Analysis of the mean neuronal trajectories revealed that the 2nd to 4th principal components (PCs), respectively, explained 8%, 5%, and 4% percent of the variance and showed significant separability between the neuronal trajectories for each stimulus (Figure 5E). The observed patterns resembled those seen with neuromast pairs: trajectories diverged in PC space at the time of the optogenetic stimulus and converged before and after.

When decoding neuromast triplets, we observed a mean decodability across neuromast combinations of 0.71 ± 0.04 (mean ± SEM, n = 12, Figure 5F) and nearly uniform values across predicted triplets. This suggests that, as stimulus complexity increases, pattern separation is slightly compromised but still robust. Population sparseness remained high as combinatorial complexity increased, whereas the number of weights recruited increased (Figure 5G and 5I). Likewise, the overlap of weights decoding different combinations remained low. In fact, the average population sparseness for pairs and triples were 0.78 ± 0.01 and 0.77 ± 0.01, respectively, whereas the mean Jaccard distance across class weights was 0.85 ± 0.005 and 0.87 ± 0.005 (mean ± SEM, n = 12, Figure 5H and 5J). These results were assonant with the need for more non-zero weights for optimal decoding of more complex stimuli (Figure 5M). For both decoders, neurons involved in the decoding process were primarily located in the lateral MON (Figure 5K and 5L).

Overall, these results suggest that small, separable ensembles of neurons in the MON can sparsely and accurately encode complex combinations of neuromast input.

### Nonlinear integration of neuromast input

Our data indicate that combinations of neuromasts are represented separately at the population level and can be decoded linearly^9,43^. We hypothesized that neuronal ensembles encoding simpler neuromast stimuli integrate nonlinearly in the MON so that a particular ensemble of neurons singularly represent combinatorial input.

To compare population activity patterns from simple and complex stimuli in the same sample, we stimulated neuromasts AP3 to AP5 individually, in pairs, or as triplets (Figure 6A). We found that when activating more neuromasts, the probability of neuronal firing of the neurons responding increased significantly (p < 0.001 for all comparisons of individual to paired stimuli and from paired to triad stimuli), whereas the number of neurons recruited increased between individual and combinatorial stimuli (p = 0.03 for comparisons between individuals and combinations, Figure 6B-C).

**Figure 6:**
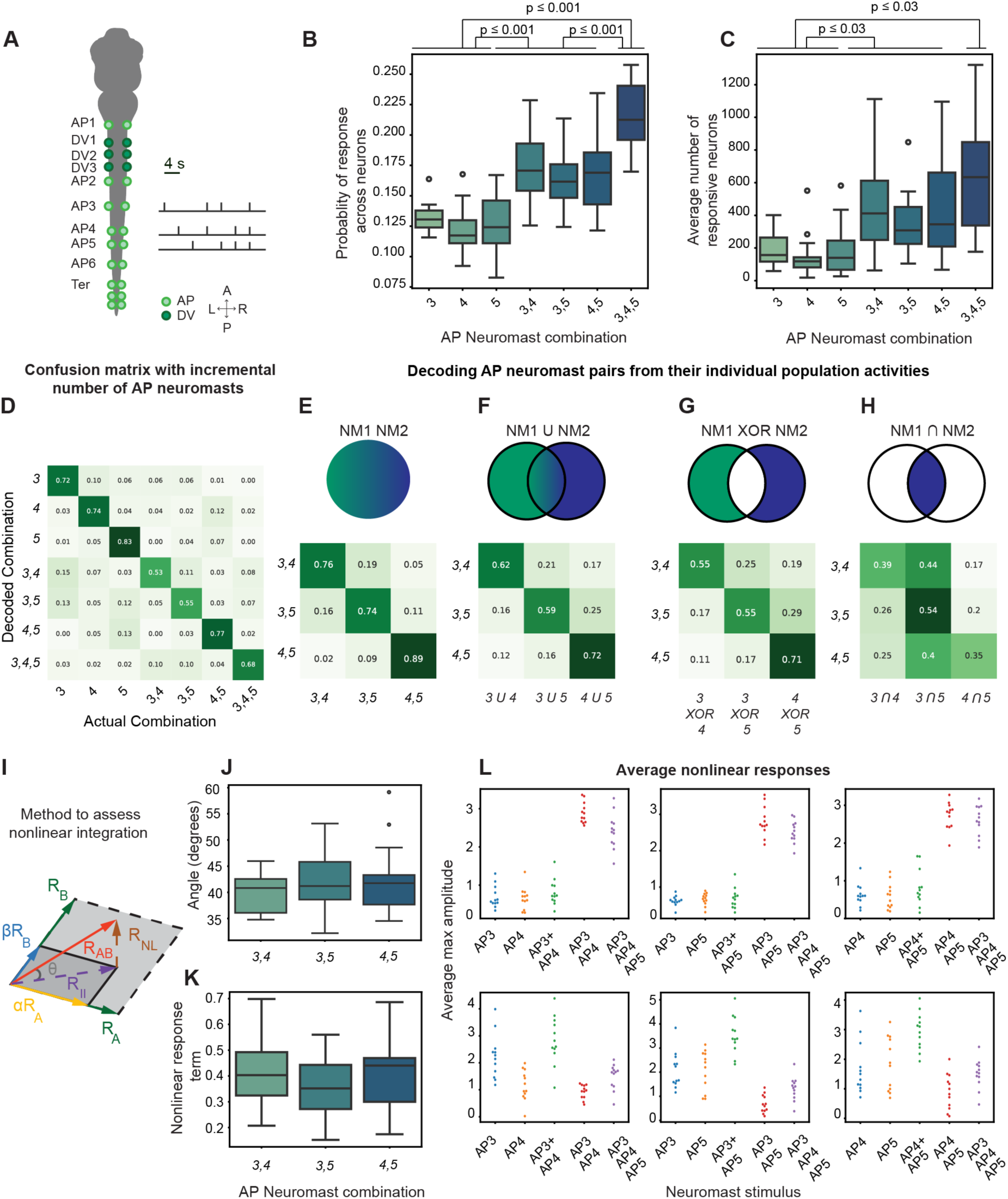
Decoding complex neuromast stimulation patterns shows nonlinear properties. **A.** A schematic diagram depicts an experiment in which we activated AP3, AP4, and AP5 either individually, in pairs, or all three neuromasts together. **B.** The average probability across fish shows that neurons fire incrementally relative to the number of neuromasts from AP3 to AP5 that are stimulated simultaneously. **C.** The average number of neurons across fish was quantified to show that more neurons are recruited when an incremental number of neuromasts from AP3 to AP5 are stimulated. For (B) and (C) statistical significance was determined using independent sample t-test. **D.** The average confusion matrix shows the decoding ability of an SVM when trained on an incremental number of neuromasts from AP3 to AP5. **E.** The average confusion matrix shows the decoding accuracy of solely pairs of neuromasts from AP3 to AP5. **F-H.** The average confusion matrices depict the decoding accuracy of the OR, XOR, and AND operators of the binarized population vectors, respectively, from individual neuromasts when used to predict the consequent neuromast pair. A schematic illustrates the vectorial analysis used to assess nonlinear integration on pairs of neuromasts, comparing it to the individual stimulation of each neuromast in the pair. The mean response of each neuron to a specific neuromast pair is compared to the optimal linear combination of the responses of each neuromast when activated individually. The difference between the mean response of the pair *(R(A,B))* and the summation of individual responses *(RA + RB)* is considered the nonlinear component of the paired stimulation *(R_NL_(A,B)).* The angle characterizing the extent of the nonlinearity can be calculated using the equation at the bottom of the panel. **J.** The values of the angle of nonlinearity across pairs of neuromasts from AP3 to AP5 show the extent of nonlinear integration when pairs of neuromasts are stimulated together. **K.** The values of the nonlinear term *(R_NL_)* across pairs of neuromasts show a tendency for supralinear integration among the population of neurons across samples. **L.** The plots show the maximum amplitude of the average calcium transient at stimulus, for all neurons with an *R_NL_* above 2 (top) and below −2 (bottom) for each sample, for the individual neuromast stimulations, the paired stimulus, and all neuromasts activated together

Overlap in neuronal ensembles constrains the number of neurons involved in sensory encoding and suggests that information is carried by interactions and dependencies between neurons. To test this, we first used our SVM to classify across all stimuli. Because in this strategy responses are binarized, assessment in decodability is purely dependent on the neuronal ensembles recruited, rather than individual changes in neuronal firing. We therefore aimed to predict the responses to paired stimuli from the responses of select populations of neurons that responded to individual neuromasts. For this, we combined the binarized vectors of individual neuromasts using AND, OR, and XOR operators (Figure 6F-H) and trained the SVM to predict the neuromast’s respective paired stimulus.

We observed a decrease in decoding accuracy when binarized vectors from individual neuromast responses were combined (Figure 6F-H), compared to a control in which we trained and predicted solely based on paired stimuli (Figure 6F, p < 0.005 for OR, p < 0.001 for XOR, and p < 0.0001 for the AND decode, n=13). However, in the OR and XOR conditions, accuracy values remained significantly above chance at values of 0.65 and 0.6, respectively (Figure 6G-H), whereas the decoding performance between these two conditions did not differ much. In contrast, when we combined binarized vectors with an AND operator, decoding on the contiguous pairs (AP3-AP4 and AP4-AP5) drops to chance levels, whereas the accuracy for the non-contiguous pair (AP3-AP5) remains close to the level observed when OR or XOR is used (Figure 6H).

To identify neurons that summate joint neuromast input nonlinearly, we used a vector-based method developed by Tejera-Leon^44^ (Figure 6I). We compared the response to simultaneous activation of neuromasts *A* and *B:* (*R(A, B)*) with the responses to each neuromast individually (*R(A, 0)* and *R(0, B)*). If *R(A, B)* is a linear combination of *R(A, 0)* and *R(0, B)*, it would lie within the plane defined by these individual responses. Any deviation from this plane indicates a nonlinearity (Figure 6J). We measured a mean angle of 43.7 ± 1.3 (n = 13) between *R(A, B)* and the *R(A,0)-R(B,0)* plane.

The mean nonlinear component (*R*_NL_*(A, B)*) was 0.43 ± 0.04 (n = 13) and remained consistent across samples and stimulus pairs, indicating a tendency of the population towards supra-linear integration (Figure 6K). Analysis of the distribution of *R*_NL_*(A, B)* values revealed that neurons with high *R*_NL_*(A, B)* values had higher calcium-transient amplitudes for paired stimuli compared to the sum of individual neuromasts, but neurons with low *R_NL_(A, B)* values showed lower amplitudes (Figure 6L). These results indicate that the integration of neuromast input involves different neuronal subpopulations that integrate individual neuromast input nonlinearly, with a slight prevalence of those showing supra-linear integration.

### Decorrelation in MON and downstream sensory areas

Using a larger number of combinations across five neuromasts, we were able to recruit neurons in the TS, OT, and medulla compared to previous experiments (Figure 7A-B). We took advantage of the neurons identified within this dataset to better compare the neuronal activity in the MON to that from other regions putatively responsive to pLL input.

**Figure 7:**
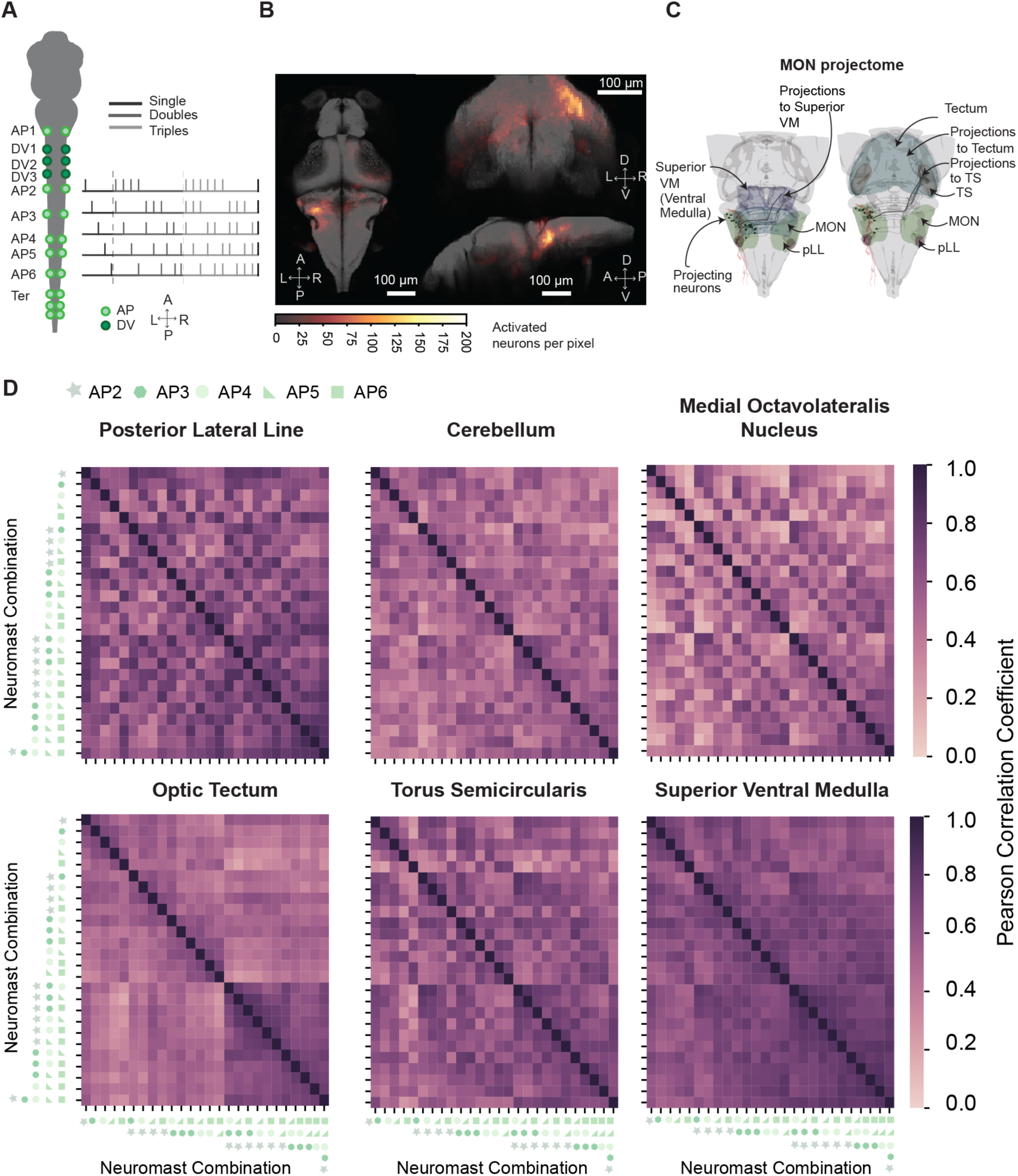
Neuronal activity in the MON is more decorrelated than that in upstream and downstream areas. **A.** In a schematic diagram of the stimulation protocol for single, double, and triads of neuromasts all 25 combinations of neuromasts of AP2 to AP6 are stimulated. In addition, all five neuromasts were also stimulated synchronously. **B.** Images illustrate the locations of responsive neurons to optogenetic neuromast stimulation across 15 fish co-registered to the MapZebrain atlas. **C.** Renderings show the projectome of the MON to the TS, OT (right), and superior medulla (left) obtained from stochastic labeling data available in the MapZebrain atlas. **D.** Inter-stimulus pairwise-correlation matrices from the mean activity of the pooled population vector across trials, of neurons in six active brain regions of the brains

In the MapZebrain Atlas, we find MON neurons projecting contralaterally to the TS, OT, and superior medulla. Projections to the TS and OT follow the medial lemniscal tract, with some MON projections traveling through the TS before turning laterally in the OT, suggesting complementary computations (Figure 7C). Conversely, neurons projecting to the superior medulla are located near where we find active neurons (Figure 7C).

We quantified similarity in firing patterns across the neuronal population within a region by calculating Pearson correlation coefficients of mean fluorescent amplitudes during stimulus presentation. The correlation matrices revealed that MON and cerebellar populations exhibited lower correlations compared to those in the pLL, TS, OT, and superior medulla, which are upstream and downstream to the MON (Figure 7D). The TS, OT, and superior medulla regions showed increased correlations for stimuli activating multiple neuromasts, whereas the MON showed higher correlations for stimuli containing the same neuromasts.

The MON exhibited slightly more decorrelation across samples compared to the pLL ganglion, OT, and superior medulla (p = 0.008, 0.04 and 0.002, n = 15 respectively), but not the TS (p = 0.28, n = 15, Figure S12). Moreover, when neurons in each region were subsampled to compensate for sampling biases between regions (p ≤ 0.001 for all regions, Figure S12). These results indicate that the MON decorrelates neuromast inputs more effectively than regions downstream, and thus is the pLL’s optimal site for pattern separation. The MON might then distribute relevant information to downstream areas like the TS, OT, and motor areas in the medulla, which could generalize the upcoming inputs for higher-order processing (Figure S13).

## Discussion

In this study, optogenetic stimulation of specific neuromasts permitted the precise activation of both individual neuromasts and groups of neuromasts along the pLL in larval zebrafish. This capability was previously limited by traditional strategies that either lacked specificity or could not apply multiplexed input^31,33,45^.

Our approach provides insights into neuromast-specific encoding. Highlighting the specificity of the optogenetic stimulus, we identified neurons with both single and mixed selectivity to neuromast input. We observed distinct spatial distribution patterns between trunk and terminal neuromasts (Figure 2D), which aligns with previous research demonstrating that trunk and terminal neuromasts exhibit different functional properties. Specifically, terminal neuromasts are innervated by larger, faster-conducting axons and tend to elicit motor responses when activated individually^16,46,47^.

We also observed a lack of spatial organization between individual neuromasts within the trunk. A lack of sensory organization is not abnormal in organisms with small brains that present constraints for a delineated spatial representation of the sensory-parameter space. For example, despite identifying cells with selectivity for various auditory properties^48–50^, in larvae, studies have not conclusively demonstrated the presence of a tonotopic organization within the MON.

Despite the absence of fine-grained topographic mapping, our data suggest that MON responses are tuned according to the neuromast’s location along the AP axis of the fish (Figure 3C). Even mixed-selective neurons follow the larva’s AP axis, suggesting convergent tuning whereby afferents with similar input locations integrate within the MON. These results indicate that sensory responses in MON neurons can be constructed from their afferent inputs in a direct feedforward manner. A similar phenomenon is observed between otolith afferents and central vestibular neurons in zebrafish^51^, which show preferential convergent input between similarly tuned afferents.

In many sensory cortices, subthreshold depolarization events occur in response to a neuron’s non-preferred input. In the zebrafish vestibular system, otolith afferents with similar tuning converge into the same vestibular neuron with a higher probability compared to afferents with more distinct tuning. We observe a similar effect in our data: mixed-selective neurons in the MON are more likely to integrate input from nearby neuromasts rather than those located further away, and decoding weights tend to exhibit high lifetime sparseness (Figure 3B-C). Therefore, this graded organization of mixed-selective input in the MON suggests that the selectivity profile of a neuron in this region is influenced by the strength of the sensory stimulus. In this context, a stronger or longer stimulation would likely result in more mixed-selective neurons, for it could generate larger post-synaptic potentials that are more likely to reach threshold and elicit spiking^52^. It remains to be determined whether stimulus discriminability is still possible under stronger stimulation.

Furthermore, the variable levels of selectivity to neuromast input are consistent with previous findings demonstrating variable complexity in MON receptive fields spanning several neuromasts. Such an arrangement helps reduce the signal-to-noise ratio between inputs while remaining flexible to various degrees of local and global hydrodynamic information. Thus, ensembles of neurons that are either single-selective or nearly so would be responsible for decoding local stimuli, such as a point source, whereas mixed neuronal populations that respond to a broad array of neighboring neuromasts would encode more global cues. This framework transcends neuromast modality, for we do not observe distinct neuronal populations responding to AP and DV neuromasts, suggesting that the pLL integrates vectorial information from a flow source in two dimensions.

Information from the pLL ganglion to the MON probably undergoes expansion recoding^2^, in which neural activity space is increased as information is projected to a larger population of neurons. This conclusion seems robust on anatomial grounds: a pLL ganglion at 6 dpf includes 60 to 80 neurons, whereas in the MON we observe in order of several hundreds of neurons responding to neuromast stimulation. Although it is possible that the observed neurons are organized into several layers of processing within the MON, an expansion of the coding subspace is still likely. Central projections of pLL neurons contain around 60 presynaptic boutons each^61^, well within the optimal expansion ratio between input and output regions for maximizing pattern separation^6,71^. Expansion recoding, accompanied by the observed decorrelation of inputs from the pLL to the MON and the high sparseness of MON responses, could be a computational strategy for encoding and separating across more complex, combinatorial neuromast input.

Expansion recoding and pattern separation are facilitated by the nonlinear integration of mixed input^43^. Given that we could decode neuromast input with a linear classifier on a binarized version of each trial’s population vector (Figures 4 and 5), nonlinearities might play a significant role in pLL neural representations^7^. Our data indicate that nonlinear integration of simultaneous activation of multiple neuromasts occurs at the population level, with relevant subpopulations of neurons summating individual neuromast input in either sub-linear or supra-linear ways. Moreover, the partial decodability we observed when predicting on the subcomponents of a neuromast pair suggests that the brain reuses some basis for representing neuromast input (Figure 6). Moreover, our results suggest that optimal decodability depends on an emergent ensemble of neurons that respond specifically when neuromasts are activated together. These neurons likely involve thresholding nonlinearities that aid in the linear separability of activity patterns^43^ (Figure 7).

Nonlinearities aid pattern separation^43,53^. However, using pseudo-simultaneous spatial input to probe the pLL only samples a limited portion of the sensory space experienced by larval zebrafish. A significant source of variation in sensory input arises from the temporal dynamics of flow stimuli^29,54^.

The separation of temporal information remains uncertain. Nonetheless, neuronal populations with nonlinear spiking might serve as coincidence detectors, potentially making them adept at processing temporal patterns^55,56^. For instance, the dendrites of MON neurons could have varying characteristics for temporal or distance integration, enabling them to detect specific delays between signals from different neuromasts^57^. This capability might generate brief periods of synchronous activity, and thereby represent complex hydrodynamic stimuli more sparsely^56^.

Our optogenetic system is well-positioned to investigate this hypothesis. The system’s fast laser-scanning capability (0.5 ms per transition) matches the maximum flow velocity, approximately 100 mm·s^-1^, experienced by zebrafish^33,54,58^. By avoiding the variability in hair bundle displacement caused by flow velocity, optogenetics additionally allows us to isolate and study the sole effects of temporal delays between neighboring neuromasts. Further experiments involving variations in temporal lags during neuromast stimulation are needed to determine whether coincidence detection is a plausible mechanism for decoding temporal flow.

Neurons in the MON that project outside this brain region typically connect to the contralateral TS and OT in the midbrain or to neurons in the superior medulla^26,59^. Our preliminary data suggest that the MON performs pattern separation of neuromast input and then relays relevant information to downstream areas based on their roles in overall brain function.

Neurons in the contralateral superior medulla likely facilitate direct sensory-motor transformations for larval responses to pLL stimuli and thus generalize the information from the MON. The role of the TS and OT in encoding hydrodynamic stimuli in 6 dpf zebrafish remains more unclear. Previous research suggests that the TS integrates multiple sensory modalities, a role that contrasts with the MON’s function in segregating sensory information^60^.

We find a few TS and OT neurons that respond to our optogenetic stimulus. Previous studies on auditory and hydrodynamic stimuli in larvae have undersampled these regions or not reported on it^25,48,50^. The TS and OT might generalize on hydrodynamic information through an ultra-sparse code^61^ or might be very selective feature detectors. The latter seems more probable. For example, in goldfish the TS and OT respond to specific types of directional stimuli with varied temporal spiking patterns^33^, and research in frogs and weakly electric fish indicates that TS neurons are generally selective to species-specific signals^62,63^. In such a case, our optogenetic stimuli might be too coarse to activate midbrain areas with high fidelity. Optogenetic stimulation of hair cells may not fully replicate the nuanced activation seen with natural stimuli, which also depends on the graded effect of water flow at each hair bundle.

Our study shows that single-neuromast optogenetics, combined with whole-brain calcium imaging, allows for precise interrogation of neuronal encoding patterns of sensory stimuli, and to define the operational limits and capabilities of a neural system. This novel technique opens the door for interrogating the encoding of complex sensory stimuli, the change of neuronal patterns under genetic or pharmacological perturbations, and the study of how neuronal encoding change during development and after nerve regeneration. Future research should also integrate our methods with global flow-based stimulation patterns, potentially within the same specimen, to connect the neuronal responses to a flow to its anatomical underpinnings. This will further solidify the pLL as a model for exploring the mechanisms behind sensory computation.

## Methods

### Lead Contact and Materials Availability

Further information and requests for resources and reagents should be directed to and will be fulfilled by the Lead Contact: Nicolas Velez-Angel. The Tg(myo6b:CoChR-GFP) line used in this study is available upon request.

### Fish Husbandry and Transgenic Lines

Experiments were performed on larval zebrafish six to seven days post-fertilization (6-7 dpf). The larvae were maintained in accordance with the standards of Rockefeller University’s Institutional Animal Care and Use Committee. They were raised in E3 medium (5 mM NaCl, 0.17 mM KCl, 0.33 mM CaCl_2_, 0.33 mM MgSO_4_, 1 mg/mL methylene blue) in an incubator kept at 28 °C with a 14/10-hour light/dark cycle.

All experiments used *Tg(elav1:H2B-jRGECO1a); Tg(myo6b:CoChR-GFP)* double transgenics on a Nacre or Casper background. The *Tg(elav1:H2B-jRGECO1a)* line was obtained from the laboratory of Misha Ahrens^37^.

The *Tg(myo6b:CoChR-GFP)* line was produced with the multisite Gateway-based Tol2 kit^64^. The CoChR-GFP construct^65^ (Addgene: 59070) was amplified through PCR with forward primer: ggggacaagtttgtacaaaaaagcaggctaagagagcgcagtcgagaggatc and reverse primer: ggggacaagtttgtacaaaaaagcaggctnngagagcgcagtcgagaggatc. The amplicon was bounded by characteristic AttB1 sites from which we performed a BP reaction to introduce the construct into a middle entry pDONR221 vector (pME COChR-GFP). We then performed an LR reaction with a 395 pDest plasmid obtained from the zebrafish Tol2 kit^64^ to insert the construct carrying myo6b:CoChR-GFP into a backbone amenable for Tol2 transgenesis and containing a cmlc2:GFP secondary marker that labels the heart. The p5E plasmid carrying the myo6b promoter was obtained from the group of Katie Kindt^66^ and the p3E-polyA was obtained as part of the original Tol2 kit^64^. Embryos were injected at the one-cell stage with 35 ng/μl of myo6b:CoChR-GFP plasmid with 120 ng/μl of Tol2 transposase mRNA. Tol2 transposase mRNA was prepared through *in vitro* transcription from a Not1-linearized pT3TS-Tol2 plasmid^36^ (Addgene ID: 31831) generated using the SP6 mMessage mMachine kit (Life Technologies) and purified by precipitation with 30 μL of 7.5M lithium chloride.

Germline transmission was identified by mating sexually mature injected progeny to Nacre or Casper adults and screened for the expression of GFP in the heart. Founders were then outcrossed to a *Tg(elav1:H2B-jRGECO1a)* line and positive larvae were raised to adulthood to establish a stable *Tg(elav1:H2B-jRGECO1a); Tg(myo6b:CoChR-GFP)* line.

### Whole-Brain Imaging

For volumetric imaging of the zebrafish brain, we constructed a light-sheet microscope based on the Swept Confocally Aligned Planar Excitation (SCAPE) design of Voleti et al. (2019)^34^. Our SCAPE microscope generates a light sheet using a 561 nm laser (Coherent, OBIS 1220123) passed through a Powell lens (Laser Line LOCP-8.9R30-1.0) and two cylindrical lenses. The light sheet is then directed by a galvanometer mirror through an illumination telescope and enters the left rim of the back aperture of a 20x water-immersion objective (1-U2B965 XLUMPlanFLx20/1.0 W, Olympus). The resulting oblique light sheet enters the sample at an angle of approximately 37° to the optical axis and is scanned across the zebrafish larva’s brain by the galvanometer.

The emitted light sheet is captured by the objective lens and travels through the conjugate plane to an acquisition telescope that magnifies the image. The light then passes through a remote scanning component consisting of two 20x objectives (1-U2B965xLUMPlanFLx20/0.95 W, Olympus) positioned at 127° to reorient and project the image onto the fast acquisition camera (Zyla-5.5-CL10-W).

The microscope setup is elevated by 31 cm from the surface of an air table, with a manual XYZ stage positioned below the imaging objective to position the sample. Our system provided a minimum resolution of 1.7 µm, 2.1 µm and 2.8 µm for the x, y, and z dimensions, respectively, and a mean resolution of 3.0 µm, 2.6 µm, 4.5 µm for the x, y, and z dimensions along the axial axis of the light sheet throughout experiments (Fig S3).

### Optogenetic Stimulation

Single neuromasts in *Tg(myo6b:CoChR-GFP); Tg(elav1:H2B-jRGECO1a)* fish were stimulated using a targeting laser positioned below the mounted sample. The apparatus includes a laser scanning system with a pair of XY galvanometers (6210, Cambridge Technology) that redirect a 488 nm beam through a 4*f* optical relay system. This system maintains one-to-one linear shift invariance of angled rays on the back-aperture of a 4x objective lens (Olympus XLFLUOR4X/340). To control the beam’s spot size, we use a slit positioned before the galvanometers. The slit is optimized to produce a laser beam diameter after the objective of approximately FWHM = 18 µm, which corresponds to 30.5 µm for 1/e^2^.

The laser stimulation system is accompanied by a fluorescent widefield microscope arm. This arsystemm consists of a 530 nm LED (ThorLabs, M530L4-C1) that overfills the 4x objective lens’s back aperture and a tube lens with a camera (FLIR: BFLY-U3-23S6M-C USB 3.0 Blackfly) that captures the fluorescence excited by the LED. This system allowed us to image neuromasts along the entire larva’s tail. To separate the optical paths for optogenetic stimulation and image acquisition, we use two dichroic mirrors.

The 4x objective lens that serves for both optogenetic stimulation and imaging is positioned below and approximately 1 mm lateral to the SCAPE’s objective lens. This arrangement prevents optical crosstalk between brain imaging and optogenetics stimulation along the tail.

Custom MATLAB software converts image coordinates into graded current inputs for the galvanometers. This arrangement allowed the precise selection of neuromast locations.

### Neuromast Identification

To label the neuromasts of 6-7 dpf *Tg(myo6b:CoChR-GFP); Tg(elav1:H2B-jRGECO1a)* larvae, we immerse larvae in 3 µM FM-4-64 for 75 s. The fluorophore enters hair cells through their transduction machinery and makes them fluoresce at 600 nm. The larvae are then screened and mounted, dorsal side up, in 1 % low-melting-point agarose in a dish with a glass coverslip bottom (Mattek: P35G-0.170).

The mounted specimen is immersed in E3 medium containing 545 µM pancuronium bromide to paralyze the fish and 2 mM L-ascorbic acid to act as an antioxidant to minimize phototoxicity. This preparation allows unobstructed access to the brain, while enabling optogenetic stimulation of the tail from the bottom through the dish’s coverslip.

In *Tg(myo6b:CoChR-GFP); Tg(elav1:H2B-jRGECO1a)* larvae, neuromast distribution typically follows the head-to-tail order described perviously^67^: AP1, DV1,DV2, DV3, AP2, AP3, AP4, AP5, AP6, and Ter. Although DV3 and AP2 occasionally swap positions, this order is accurate more than 90 % of the time. DV3, having fewer hair cells, appears dimmer during screening and can easily be discriminated from other neuromasts. Moreover, AP6, which is sometimes referred to separately from the terminal neuromasts, was occasionally absent.

When there was uncertainty regarding the orientation (AP or DV) and the order of the neuromasts targeted, larvae were fixed, stained with phalloidin, visualized at high magnification with a confocal microscope, and cross-referenced to the tail’s image obtained during the experiment.

### Optogenetic laser power and scattering measurements

Optogenetic laser power is quantified with a Thorlabs Photodiode Power Sensor (Model: S120VC). For most experiments, the laser is operated at 41 mW·mm^-2^, which corresponds to 0.03 mW at the specimen, as calculated for a spot diameter of 30.6 µm as defined by the PSF’s 1/e^2^ value. That power density resembles the levels reported for one-photon optogenetics in a neuronal culture system and in *ex vivo* preparations^68^. Power titration experiments were also conducted to ensure that 41 mW·mm^-2^ activated neuromasts effectively, by activating three subsequent neuromasts in the trunk of the fish individually.

Scattering measurements were performed by imaging the scattering profile of the 488 nm laser beam as it interacted with the zebrafish skin. The laser power density used for this characterization was of 41 mW·mm^-2^. We measured a cross-section of the laser’s back-scattering profile across several points along the AP axis of the zebrafish. These profiles were fitted with a sum of two Gaussian functions, one representing the target intensity and the other accounting for the tails of the distribution due to light scattering from tissue^69^. The spot size of the scattered beam, characterized by its FWHM, was found to be on average, 12.34 ± 1.32 µm (mean ± SEM, n = 113) and it did not vary significantly along the larva’s trunk. That of the second Gaussian, denoting the long-tailed spread of the scattering was 41.5 ± 1.37 (mean, SEM, n = 113). We compared the intensity of the second Gaussian at 100 µm from its center, a conservative minimum for the distance between neuromasts, to the thresholds for neuromast laser power activation described above. We found that laser power 100 µm from the target was far lower than the minimum power needed to elicit depolarization in a neuromasts.

### Parameters of optogenetic stimulation

Each stimulus had a total duration of 50 ms. When stimulating an individual neuromast, this period consisted of discrete light exposures of 1 ms pulses at least every 5 ms. When activating a combination of neuromasts, each neuromast was exposed to light for 1 ms at a time, with the laser transitioning between neuromasts in an oscillatory fashion with a minimal elapsed time of no more than 0.5 ms. To equate to the stimulation frequency in individual neuromasts, we controlled the transition between neuromasts such that, on average, each neuromast in the combination was repeatedly stimulated at an average frequency of 200 Hz. For pairs of neuromasts, this resulted in a transition time of 1.5 ms, whereas triplets had a transition time of 0.67 ms. We consider this stimulation nearly simultaneous, for the time delay between transitions is below the average rise time of CoChR-GFP activity^36^ (4-6 ms), which allows the integration of information across neuromasts.

For the experiments depicted in Figure 2 and Figure 3, each neuromast was stimulated once per minute. This approach ensured consistency in stimulation despite the varying number of neuromasts that were accessible in each experiment. Consequently, the inter-stimulus intervals between neuromasts within a trial ranged from 6 s to 10 s, with a limit of 20 trials per experiment.

For experiments involving a predefined set of neuromasts (Figure 4 and subsequent figures), the interstimulus interval was 4 s. The number of repeat trials depended on the number of neuromast combinations used in each experiment for recordings that totaled 26 min.

In addition, for experiments focused on decoding neuromast stimuli (Figs. 4-7), the order of stimulation for each trial was randomized.

### Image processing

Volumetric calcium imaging data were acquired at 2.18 volumes per second. The dimensions of the volume were X:1008 µm x Y:500 µm x Z:316 µm, which encompassed the larva’s entire brain. Sweeping the light sheet along the X-axis at intervals of 2 µm resulted in 250 slices. The data were transposed and every three slices in the axial dimension were averaged. Each slice was then filtered and motion-corrected by non-rigid motion correction (NoRMCorre)^70^. Fluorescence traces for individual cells were extracted using the constrained non-negative matrix factorization (CNMF) option from CaImAn^71^. The autoregressive model order in CaImAn was set to one, and the gSig parameter was set to two for neuronal nuclei with an average radius of 5 µm. ROIs identified by the CNMF algorithm were then detrended within CaImAn. Finally, ROIs were collated across planes to remove doublets appearing across nearby axial slices. Doublets were assessed as any ROIs within five pixels of each other and on contiguous planes, with a temporal correlation coefficient above 0.85. Samples that did not segment accurately were discarded.

### Registration of zebrafish brain volumes

Registration of brain volumes was performed following the protocol and parameters outlined^72^. Advanced normalization tools (ANTs, https://github.com/ANTsX/ANTs, ^73^) was used to create a template volume from all experiments in a dataset. Symmetric diffeomorphic normalization^74^ was then applied to register this template to the H2B-GCaMP reference brain from the MapZebrain Atlas^30^. High-resolution volumes acquired before each experiment were used to construct the template. The resulting warps from the registration to the atlas were applied to the XYZ coordinates of the ROIs and assigned to the annotated regions of the MapZeBrain Atlas. Samples that did not register accurately were discarded.

### Selection of neuromast-responsive neurons

To identify statistically significant neural responses, we adopted the approach described by Demas et al.^75^. For both single and combinatorial stimuli, we identified neuromast-specific responsive neurons by correlating the time series of each neuron’s calcium imaging with a stimulus vector that represented the times at which a neuromast was stimulated. The stimulus vector was generated by convolving a binary time series —matching the sampling rate of the acquired imaging data— in which a“1” indicated the time-point of stimulation and “0” elsewhere, with a double exponential kernel designed to model the decay kinetics of the calcium indicator. For our study with H2B:jRGECO1a, we estimated the decay constant^76^ of the indicator to be less than 3 s. All correlations were computed using Pearson’s correlation coefficient, and traces were min-max normalized before calculating the correlations.

To assess statistical significance, we performed null-hypothesis testing by creating convolved time series with randomly shuffled stimuli and correlating them with all segmented neurons. The number of randomly shuffled stimuli matched the number of times a neuromast was activated. This process was repeated 500 times to establish the null distribution. The distribution of null R^2^ values was fitted to a normal distribution, and neurons were considered significantly active if their correlation was at least 2.5 standard deviations above the mean of the null distribution.

### Topography analysis

To visualize the neurons activated by single neuromast input and study the spatial organization of neurons responsive to different neuromasts along the anterior-posterior axis, we first selected neurons that were correlated to solely one neuromast and not others, as determined by the method described earlier. To map the distribution of responsive ROIs in the MON, we used the spatial positions of each group of neuromast-selective ROIs as input the computation of a one-dimensional kernel density estimation (KDE).

Comparisons between the distributions of XYZ coordinates of ROIs responsive to each neuromast were conducted by calculating the pairwise Kullback-Leibler divergence (KLD) between each set of registered XYZ coordinates in the pooled dataset, except for AP1, which offered very few examples. Because the KLD metric is asymmetric, we report the average between the metric the normal and flipped comparisons between the two distributions.

We additionally assessed cluster homogeneity by calculating the relative same-class probability for each ROI and its neighboring ROIs within a specific radius. For this analysis, we used the PyNNDescent package in Python for accurate nearest-neighbor search, sampling up to 2,000 neighbors per iteration and constructing a connectivity graph in which nodes represent ROIs and edges represent distance values. We computed the probability of encountering neurons responding to the same neuromast for a given distance and controlled for sampling bias by normalizing that probability value by the overall probability of finding a neuron responsive to a particular neuromast.

### Analysis on single- and mixed-selectivity populations of neurons

To evaluate selectivity, we first averaged neuronal activity across trials and z-scored. A neuron was classified as responsive to a particular neuromast if its mean amplitude during optogenetic stimulation was at least two standard deviations above the mean of a median filtered baseline (window of four frames) for a period of one frame before the stimulus and three frames after stimulation (a total of 1.8 s).

Neurons were categorized as single-selective if they met this criterion for only one neuromast among those targeted during the experiment. Conversely, mixed-selective neurons were those responsive to more than one neuromast. From this analysis, we calculated the conditional probability that a neuron selective to neuromast A was also selective to neuromast B across all possible neuromast combinations. This was determined by dividing the number of neurons responsive to both neuromast A and neuromast B by the total number of neurons selective to neuromast A.

For the part of the study in which we assessed whether mixed-selective neurons carry information about neuromast identity, we selected the mixed-selective population by identifying neuromast-responsive neurons across combinations of single-neuromast stimulus trains, up to the point at which all optogenetic stimuli were represented. As described in the section above, we identified stimuli-responsive neurons by correlating each neuron with each stimulus train, obtained the Pearson correlation coefficient, and then compared this value to a null distribution. To isolate the mixed-selective population, we excluded any neurons that showed a significant correlation with the stimulus train of the stimulation of only one individual neuromast. Such neurons corresponded to the dataset used for previous analyses.

### Analysis of neuronal trajectories in PC space

To visualize neuronal trajectories, we analyzed mean traces from an 8-second peri-stimulus window (4 s before and after each optogenetic stimulus), resulting in an *N* × *t* matrix, where *N* is the number of neurons responsive to neuromast input and *t* is the time window. We concatenated these traces along the *t*_-_axis and *z*-scored them. We then performed SVD and projected the data onto the principal components determined by the matrix’s smallest dimension. We repeated this analysis for both individual samples and the pooled neuron population, obtaining similar results.

The first principal component (PC 0) did not differentiate between stimuli, but PCs 1, 2, and 3 did. Therefore, we plotted the mean trajectories in 3D space for PCs 1, 2, and 3 in three dimensions smoothing each trajectory visualization with a Gaussian kernel of one.

### Decoding analysis

To determine if population activity can discriminate between different neuromast stimuli, we trained multiclass support vector machine (SVM) classifiers with a linear kernel using the LinearSVM implementation from the scikit-learn toolbox in Python.

We conducted the decoding analysis on the subset of neuromast-responsive neurons identified through the correlation analysis described above. For decoding stimuli that targeted individual neuromasts, we only included samples with more than 100 identified neurons across stimulus categories. For combinations of neuromasts, we only used samples with more than 200 neurons identified across neuromasts.

For each sample, we applied one-shot encoding to the time trace values of each responsive neuron during the stimulus period. As for the selectivity analysis, if the peak amplitude activity within four frames (1.8 s) from the time of stimulation was two standard deviations above the baseline’s mean, we labeled it as one for that neuron. Otherwise, it was labeled as zero. This transformation resulted in a tensor of binary values with dimensions *N* x *T* x *S*, in which N denotes the number of neurons, *T* denotes the number of trials, and *S* denotes the stimulus categories.

We trained the classifier using 80 % of the binarized population vectors per stimulus and tested it on the remaining 20 %. Both training and testing sets were sampled randomly. The training set underwent ten-fold cross-validation, in which the regularization hyperparameter of the SVM was iterated from 0.1 to 10. We selected the weights that provided the highest accuracy in comparison to the annotated neuromast input labels from each held-out validation set. The weights across validation sets were averaged to represent the optimal hyperplane for separation.

### Analysis of SVM weights

We analyzed the weights of the decoder by measuring its population and lifetime sparseness. Lifetime sparseness measures the degree to which a single neuron’s weights are distributed across different stimulus categories. A lifetime sparseness value of 0 indicates that a neuron contributes equally to all stimulus categories, whereas a value of 1 indicates that it contributes exclusively to one stimulus category.

Population sparseness quantifies how sparsely the ensemble of weights contributes to decoding a specific stimulus category. A population sparseness value of 1 suggests that most neurons are involved in the decoding effort, whereas a value of 0 indicates that only a few neurons are responsible for carrying the information about the stimulus category.

We measured sparseness using Hoyer’s method^77^, which is defined as follows:

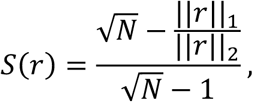

in which *N* is the number of distinct stimuli and *r* is the value of the weights per class per neuron. By quantifying the degree of sparseness in the distribution of weights for each neuron and across the population, this equation reflects how effectively the neuronal responses are utilized for distinguishing between different stimuli.

To assess the overlap between weights across categories, we obtained the indices of non-zero weights for each weight category vector, creating a vector of indices for each category. We then calculated the Jaccard dissimilarity index between these vectors. If the Jaccard dissimilarity index approaches 1, then the vectors of indices have no overlap with one another, whereas if the index approaches 0 it means the vector of indices across categories remains the same.

### Analysis of nonlinear summation

In experiments involving both single and double neuromast stimulation within the same sample, we compared the population vector of responsive neurons for each neuromast pair with the population vectors for each neuromast component^44^. This comparison was performed on mean responses across trials to mitigate the effects of trial-to-trial variability. The mean responses for each stimulus were consolidated into a single population vector by taking the maximum value within four frames (1.8 s) following the optogenetic stimulus. To address potential issues such as photobleaching of the calcium indicator’s activated state in some neurons and potential habituation to repeated stimuli, we applied min-max normalization to the values of each trace.

Given neuromasts *A* and *B*, we measured the angle between the dual neuromast response *R(A,B)* and the plane defined by the responses to the individual neuromasts *R(0,B)*. This was achieved by first minimizing the L2 norm of the difference between *R(A,B)* and the linear combination α *∗ R(A,0) +* β *∗ R(0,B)*. The minimized error, denoted as *R*_NL_*(A,B)*, reflects the degree of nonlinearity in the response to the paired stimulus. The minimized equation is then the following:

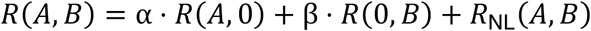

A positive *R_NL_(A, B)* indicates supra-linear summation, in which the response to the combined stimuli is greater than the sum of individual responses. Conversely, a negative *R*_NL_*(A,B)* signifies sub-linear summation, in which the response to the combined stimuli is less than expected.

The angle between *R(A,B)* and the plane spanned by *R(A,0)* and *R(0,B)* was calculated using the formula

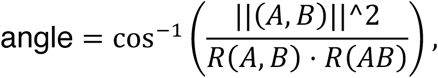

which measures the extent of nonlinear summation in neuronal populations responding to individual versus paired stimuli. Values close to 0° degrees indicate negligible nonlinearity, whereas values approaching 90° (π/2) suggest that *R(A,B)* resides orthogonally to the plane defined by *R(A,0)* and *R(0,B)*. We validated this approach by ensuring that the alpha and beta coefficients in the minimization procedure were bounded between −1 and 1 and were consistent across different combinations. This validation was crucial to confirm that our measurements were reliable and not artifacts of the noise structure.

To further validate our results, we performed the same calculations for non-correspondent pairs of neuromasts to assess the robustness of our measurements. We also compared our results to a neuronal matrix of average trial responses in which the population structure was scrambled. In both cases we compared the angle values from these null-model pairs with those from the actual neuromast pair comparisons and their corresponding individual counterparts. Although the null models did not exhibit angles close to orthogonality—as expected—they showed higher angles compared to the actual neuromast pair comparisons. This confirmed that the differences we observed were meaningful and not due to noise.

### Pattern decorrelation across individual and combinatorial neuromast input

Population vectors of neuronal activity were determined by taking the mean amplitude for each trial averaged neuromast responsive neuron within a 1.8 s stimulus window^41^. Using the region annotations provided by the MapZebrain Atlas^30^, we segmented the ROIs from all relevant anatomical areas. The Pearson correlation coefficient was computed between the mean population vectors for each pair of stimulus conditions to yield in a correlation matrix for each brain region. To account for potential biases in these measurements per experimental sample, we performed the same analysis for each individual larva and then averaged the Pearson correlation coefficients for each pair of conditions.

To control potential sampling biases across brain regions, we also randomly subsampled 200 neurons from each region and repeated the analysis as described. This process was iterated 1000 times, and the average Pearson correlation coefficient per region was calculated for each subsample instance.

## Acknowledgments

The authors wish to thank the members of the Hudspeth lab for valuable advice and suggestions on the manuscript; Alipasha Vaziri, Priya Rajasethupathy, Vanessa Ruta and Marcello Magnasco for useful discussions; Jeffrey Demas, Tobias Noebauer, and Sanjeewa Abeytunge for help with optogenetic stimulation; and Elizabeth Hillman, Citlali Perez Campos, and Wenze Li for help with building the SCAPE microscope. Thanks to Natalia Velez for feedback on graphical design. The dissemination of the SCAPE microscope was supported by the NIH BRAIN grants UF1 NS108213 and U01NS094296. The rest of the study was supported by Howard Hughes Medical Institute.

## Author contributions

NVA, AJ and AJH were involved in the conceptualization of the work and wrote the manuscript. NVA and BF did experiments with help of YV. NVA, SL, BF, CR and HB contributed to data analysis. All authors edited the manuscript.

