## Supplementary Information for "Optogenetic interrogation of the lateral-line sensory system reveals mechanisms of pattern separation in the zebrafish brain"

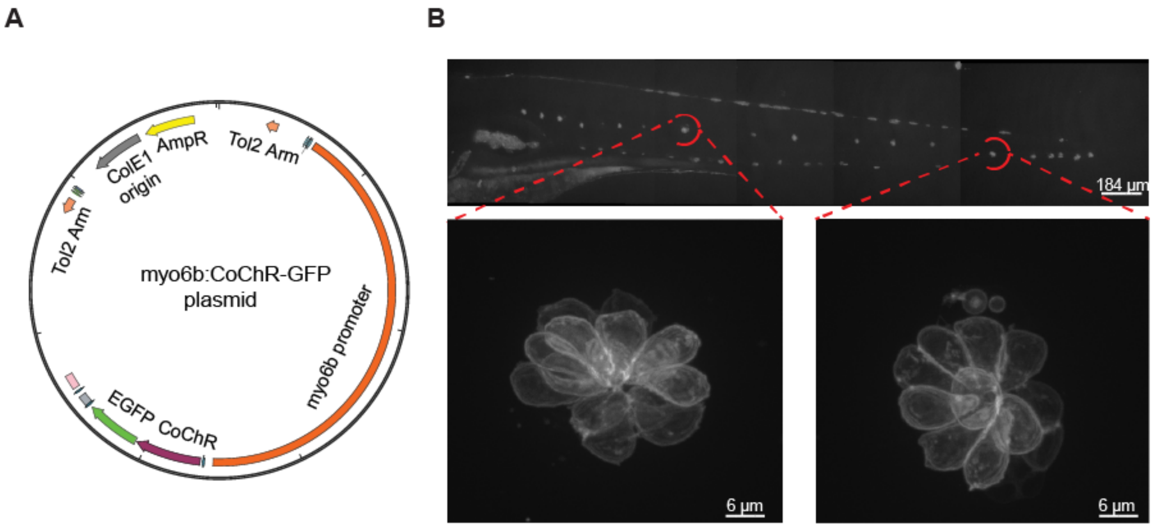


**Figure S1: Characterization of *Tg(myo6b:CoChR-GFP)* fish: A.** A map of the plasmid used for the transgenesis of the *Tg(myo6b:CoChR-GFP)* line. The plasmid carries an approximately 6 kb fragment of the *myo6b* promoter, which targets expression to hair cells; a copy of the CoChR-GFP transgene; and Tol2 arms for Tol2 transposition into the zebrafish genome. **B.** The top image of a 6 dpf *Tg(myo6b:CoChR-GFP)* fish shows expression of CoChR-GFP in the neuromasts of the pLL. The two example images at the bottom show the expression pattern of CoChR-GFP within a neuromast. As expected CoChR-GFP localizes to the plasma membranes of hair cells.

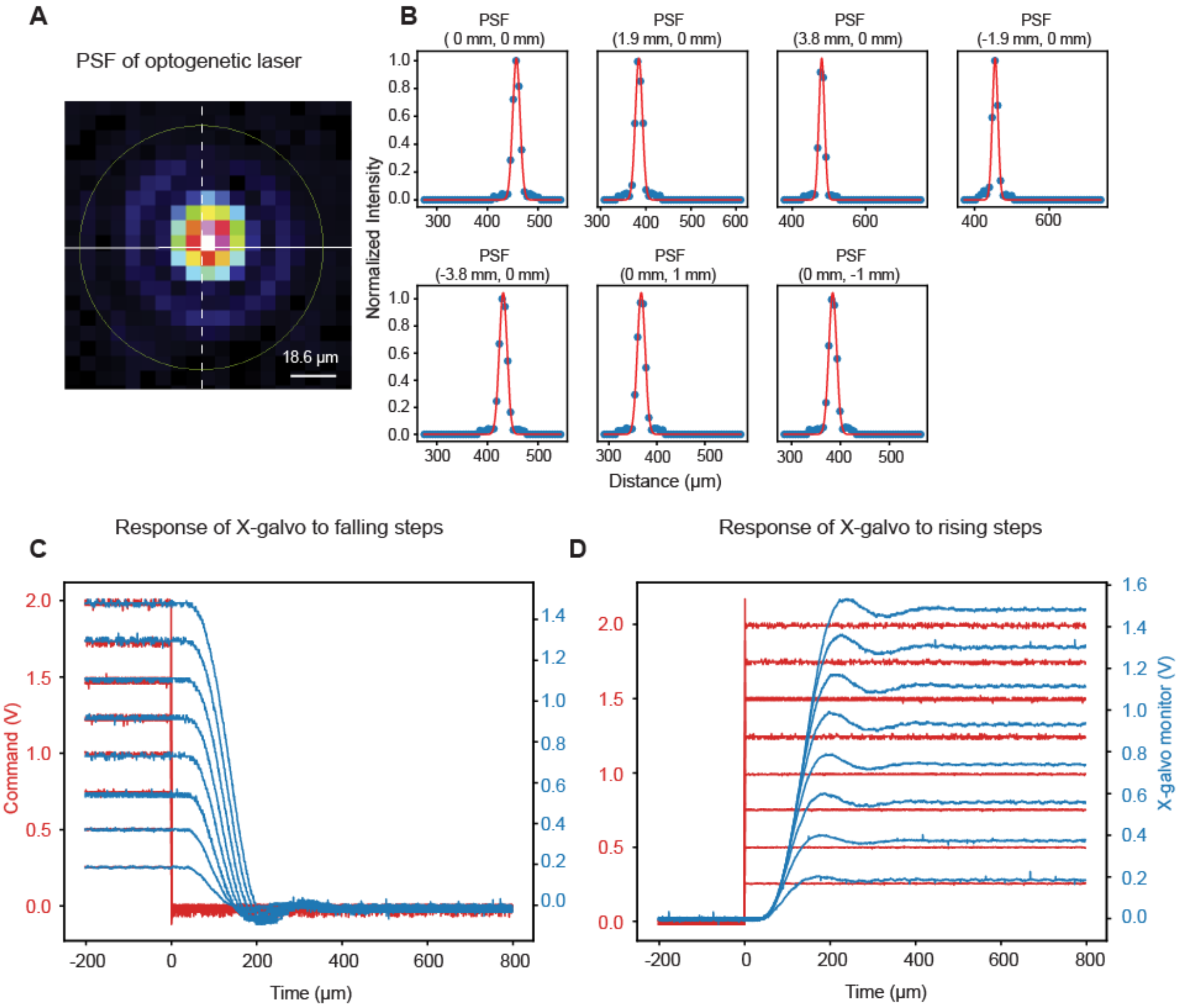


**Figure S2: Characterization of the optogenetic laser system: A.** A beam profiler provided an image of the point-spread function (PSF) of the optogenetic beam at its focus. The measured full width at half-height is around 18 µm, a diameter slightly smaller than that of the area occupied by hair cells in an average neuromast. **B.** The plots show the profile of the optogenetic beam as it is displaced from the center to the extremes of the field of view. As expected with a diffraction-limited 4f system, the beam profile does not vary at the X and Y extremes of the field of view. **C-D.** Two plots portray the equilibration dynamics of the X galvanometer mirror to falling and rising voltage steps. This is the voltage range needed to relocate the laser from one edge of the 3.8 mm field of view to the opposite edge. The blue traces show that the movement in response to the commanded voltage reaches a steady state in 500 µs, which permits nearly simultaneous activation of neuromasts.

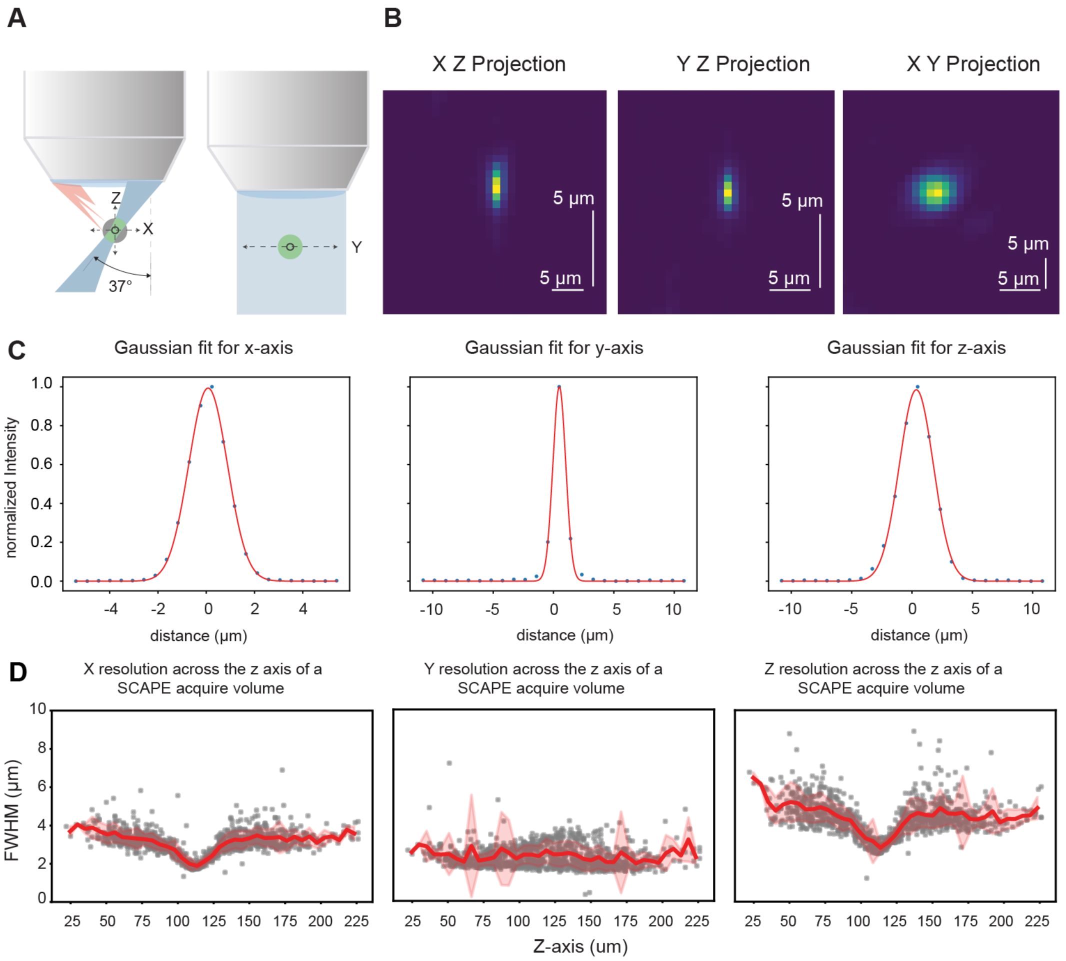


**Figure S3: Characterization of the optical resolution of the SCAPE system over its field of view: A.** An illustration of the axes along which resolution was measured by imaging 1 µm beads. The right image shows a frontal view of the microscope objective, whereas the left image shows a side view. **B.** The images display a representative example of a 1 µm bead at the center of the field of view (within the beam waist of the SCAPE's light sheet). **C.** The plots show the X, Y, and Z cross-sections of the bead in B. fitted with Gaussian profiles. **D.** The plots provide the resolution of the system along the Z axes. The variation in resolution is dependent on the beam's waist: farther from the waist, the light sheet widens and thus the resolution is compromised.

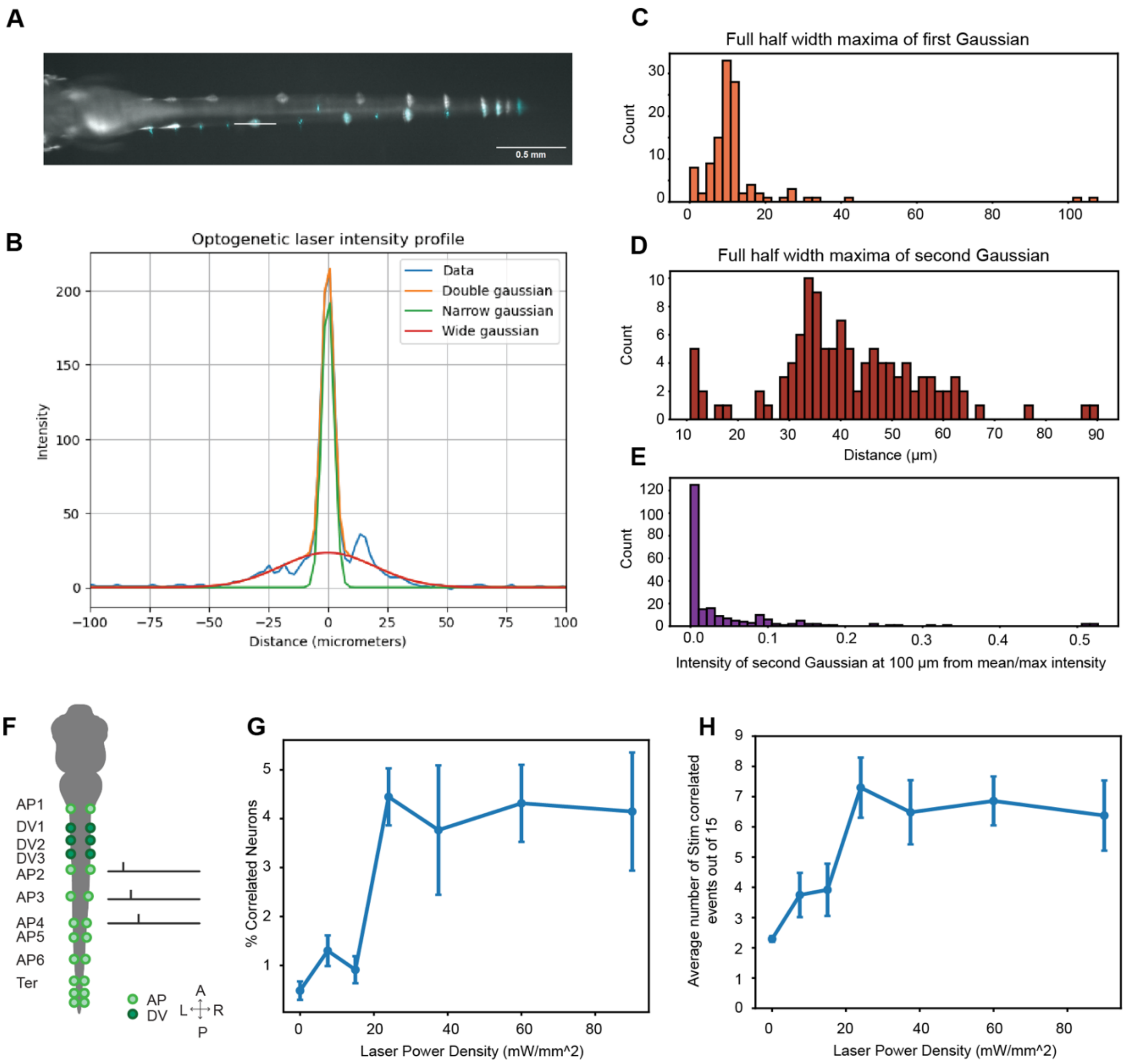


**Figure S4: Scattering and crosstalk between neuromasts during optogenetic stimulation: A.** In order to measure the spread of the laser beam when scattering with the fish's tissue, an image of the tail of a zebrafish larva is overlayed with the scattering profiles from the laser beam at different locations along the larva's AP axis. **B.** The plot displays an example of a laser's scattering profile cross-section fitted with a double Gaussian function. One of the functions fits the signal's power and the other one fits the tails of the scattering profile. **C-D.** Two histograms show the variations of the full-width half-maxima of the first (C) and second (D) fitted Gaussian functions along the AP body axis of each of seven larvae. **E.** The distribution of normalized intensity values of the second function 100 μm from the curve's peak indicates that at this distance, there is negligible light from the targeting laser. **F.** A schematic illustrates the activation of three neuromasts individually with an inter-stimulus interval of 10 s. Different sets of samples (n = 4-6) were activated for each power density tested. **G.** The power density used for most experiments was confirmed by quantifying the number of correlated neurons to single neuromast stimulus across the interrogated power densities and **H.** The average number of events elicited by single neuromast stimulation, out of a total of 15, across the range of power densities.

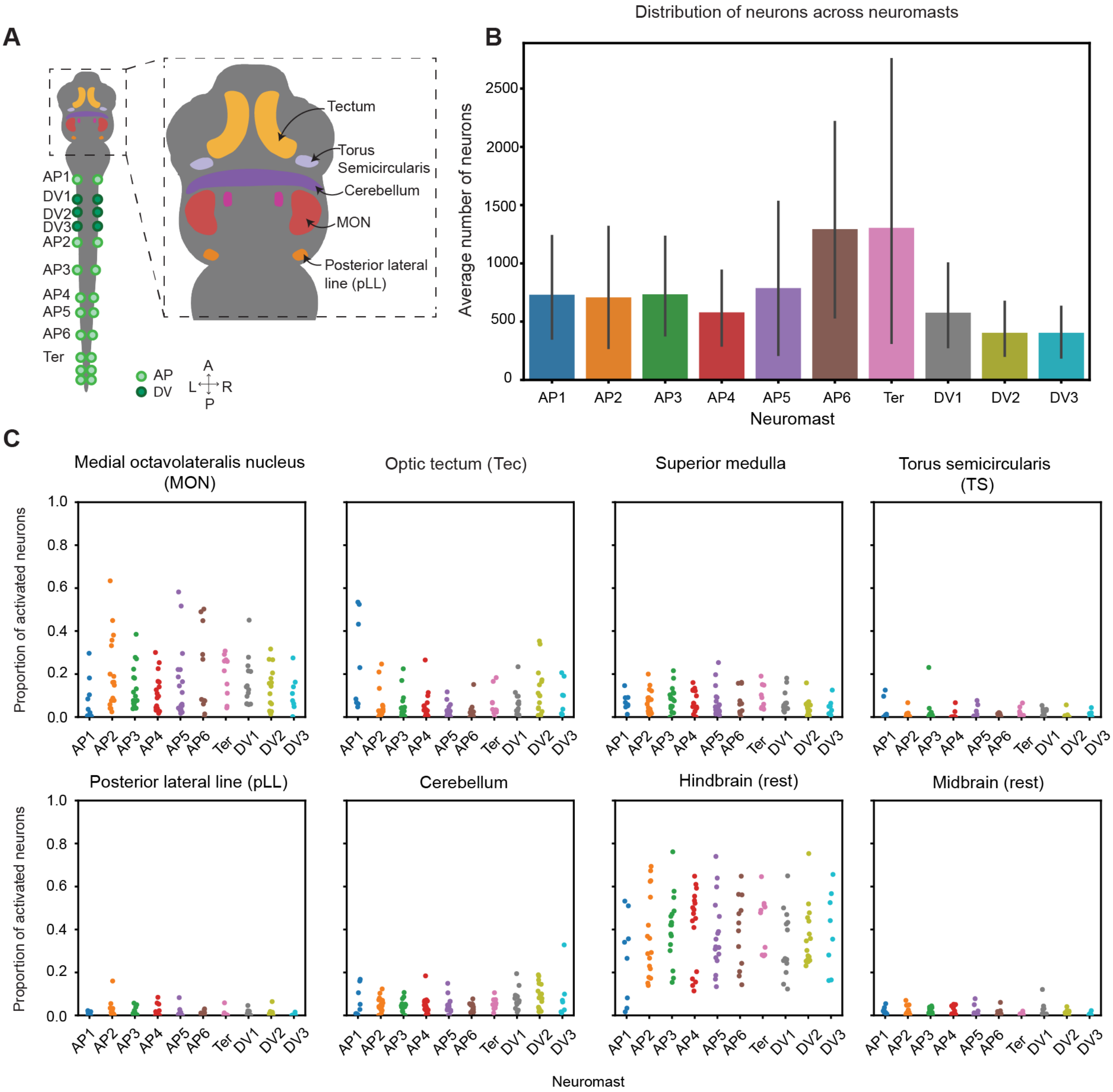


**Figure S5: Quantification of the number of neurons activated when individual neuromasts are activated with optogenetics. A.** A schematic depicts the areas of the zebrafish brain that we expect to be activated by the optogenetic stimulation of neuromasts. **B**. A barplot quantifies the number of neurons, on average, that are activated by the optogenetic stimulation of each neuromast. Neurons in the brain are active equally across neuromasts. The plot depicts the mean across samples and the error bars correspond to the standard error of the mean (SEM). **C.** Swarmplots show the same quantification in B. but separated across eight regions within the zebrafish brain. We find prolific activation in the medial octavolateralis nucleus (MON) as well as the rest of the hindbrain, while more mild activation in the other six regions. The number of neurons is normalized by the total number of neurons that significantly correlated to that neuromast across samples.

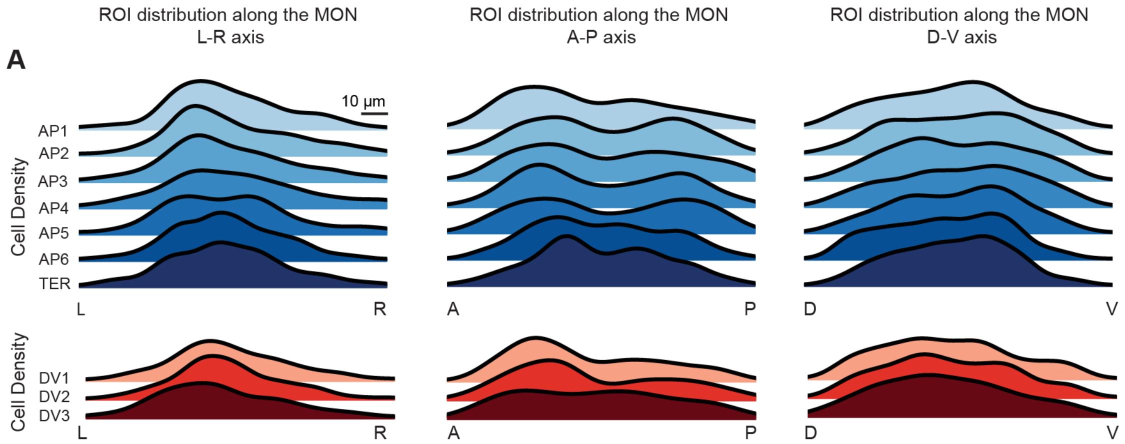


**Figure S6: Spatial distributions of neurons in the MON in response to each neuromast. A.** The spatial densities of neurons responsive to each neuromast across the MON are highly overlapping. In order: the following axes are represented: the lateral axis (right), anterior-posterior axis (center), and dorsalventral axis (left), depicted as a kernel density plot. Densities corresponding to AP neuromasts are shown in blue and to DV neuromasts in red. Axis directionality follows the same conventions as in B.

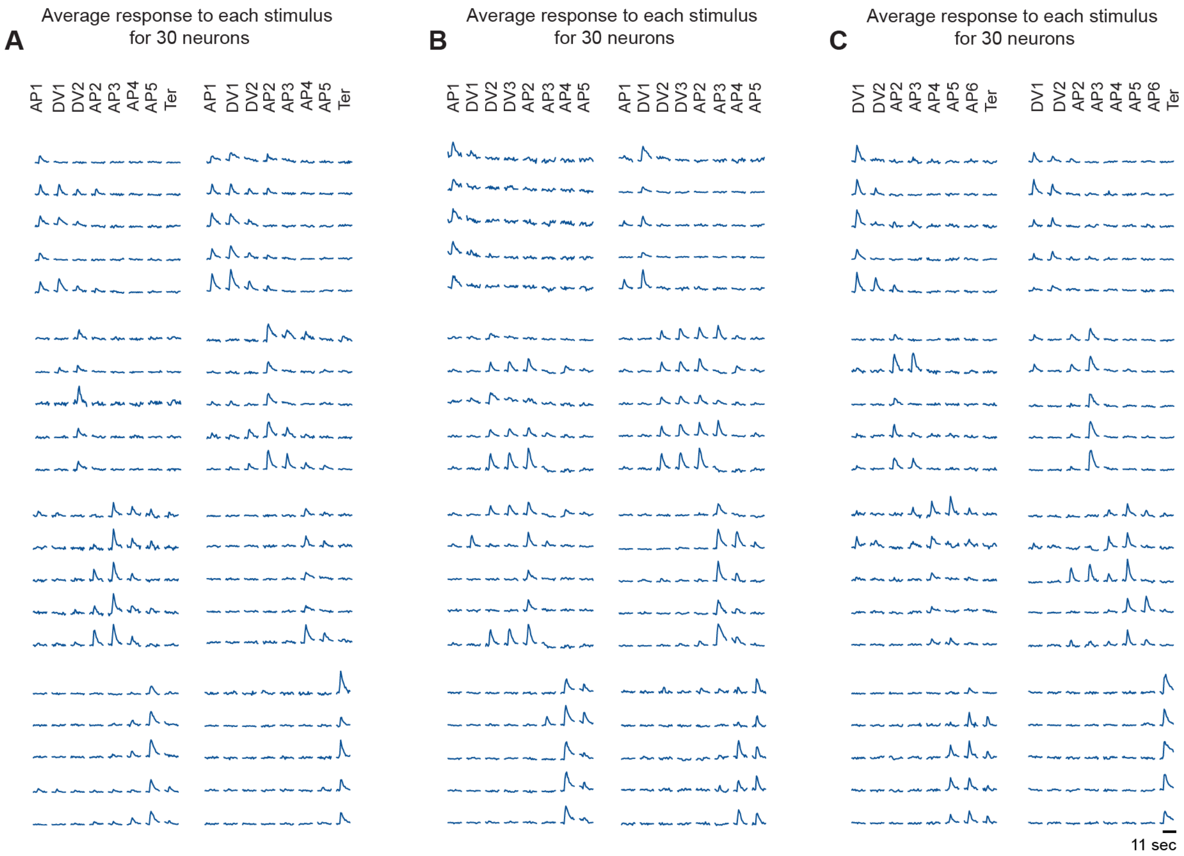


**Figure S7: Example mean traces for the activation of individual neuromasts. A, B, and C.** In all three cases, the figure depicts the z-scored mean activity of five example neurons with a correlated response to the optogenetic stimulation from each of eight neuromasts targeted individually during an experiment. Examples in each block, for each fish, reveal the presence of single and mixed selective neurons following the body axis of the zebrafish larva in a similar fashion to Figure 3b.

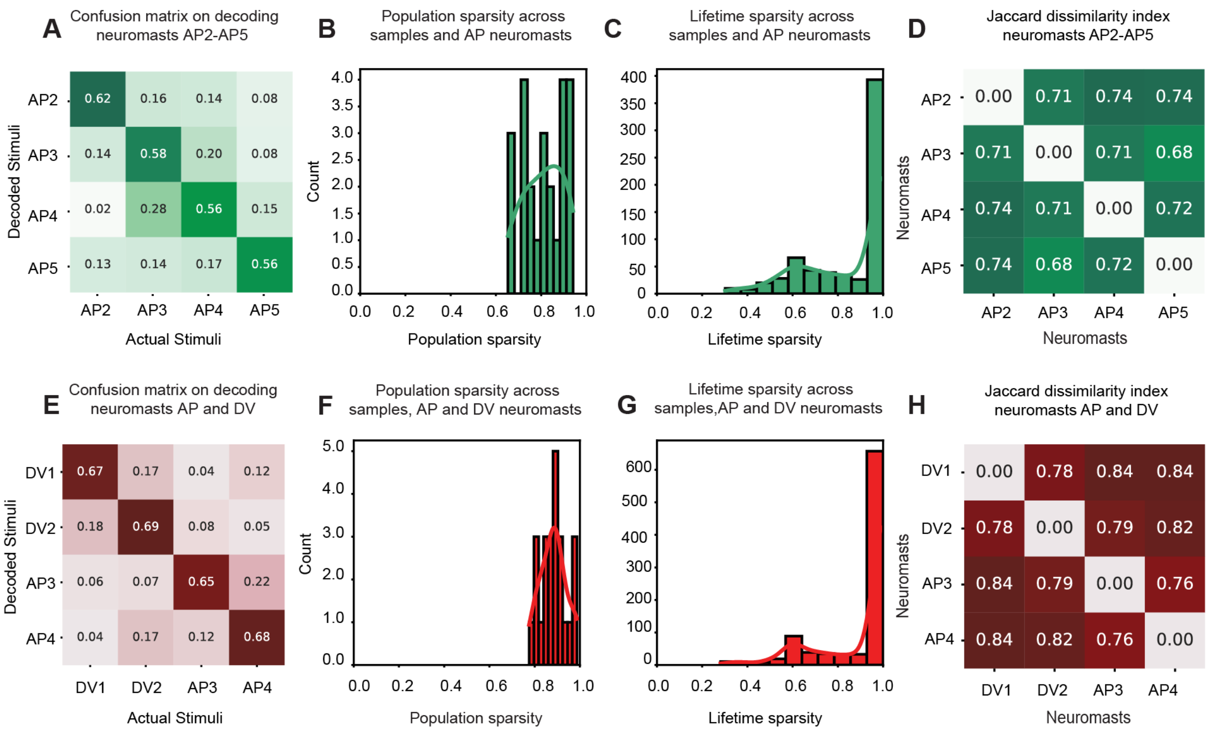


**Figure S8: Decodability of individual neuromasts considering a mixed selective population of neurons in the zebrafish brain. A.** The confusion matrix for the classification of neuromast input between anterior-posterior (AP) neuromasts (AP2-AP5) shows lower decodability when predicting from a mixed selective population of neurons compared to when only single selective neurons are included. **B.** The sparsity of the weights across the rows for the SVM classifier in A. illustrates the sparsity in the mixed populations of neurons decoding neuromast identity. **C.** The sparsity of the weights across the columns for the SVM classifier in A. indicates the tendency for weights to be involved in predicting one or multiple neuromast categories. **D.** The Jaccard dissimilarity index of the non-zero weights across neuromast classes for the SVM classifier in A. summarizes how weights are distributed across neuromast categories. **E.** The confusion matrix for the classification of neuromast input between dorsal-ventral (DV) and AP neuromasts (DV1, DV2, AP3, AP4) shows lower decodability when predicting from a mixed selective population of neurons compared to when only single selective neurons are included. **F.** The sparsity of the weights across the rows for the SVM classifier in E. illustrates the sparsity in the mixed populations of neurons decoding neuromast identity. **G.** The sparsity of the weights across the columns for the SVM classifier in E. indicates the tendency for weights to be involved in predicting one or multiple neuromast categories. **H.** The Jaccard dissimilarity index of the non-zero weights across neuromast classes for the SVM classifier in E. summarizes how weights are distributed across neuromasts.

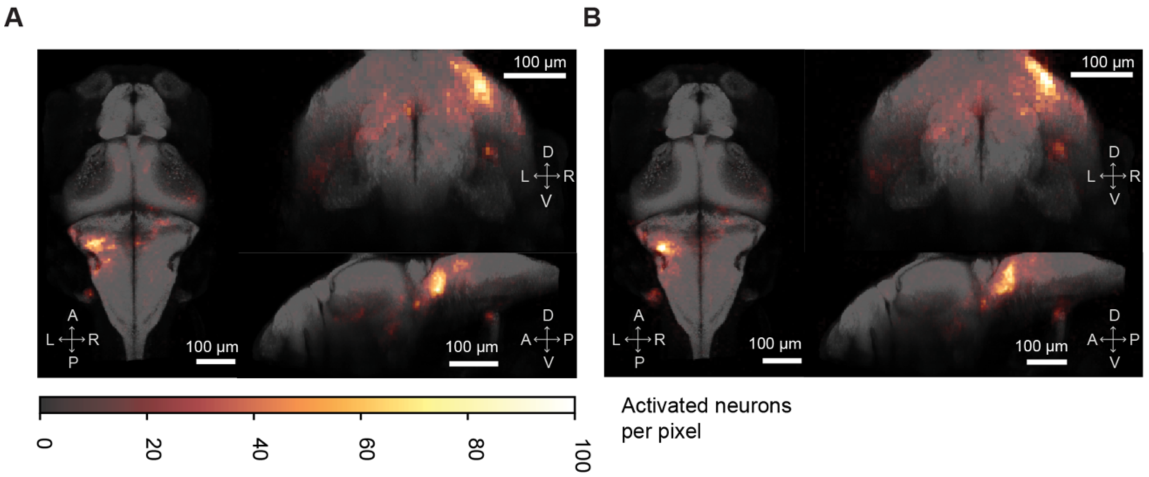


**Figure S9: Spatial location of neurons responding to combinatorial stimuli. A and B.** The locations of neurons responsive to stimulation of pairs of neuromasts (A) and neuromast triplets (B) in 12 fish co-registered to the MapZebrain atlas. Images show activity primarily in the ipsilateral MON but also in the superior medulla, the TS, the OT, and the ipsilateral cerebellar crest. For both A and B the coordinate axes is as follows. Left panel: Horizontal maximal projection. A: Anterior; P, Posterior; L, left; R, right. Right upper panel: Coronal maximal projection. D, Dorsal; V, Ventral; L, left; R, right. Lower right panel: Sagittal maximal projection. D, Dorsal; V, Ventral, A: Anterior; P: Posterior. The scale bar for all images is 100 µm.

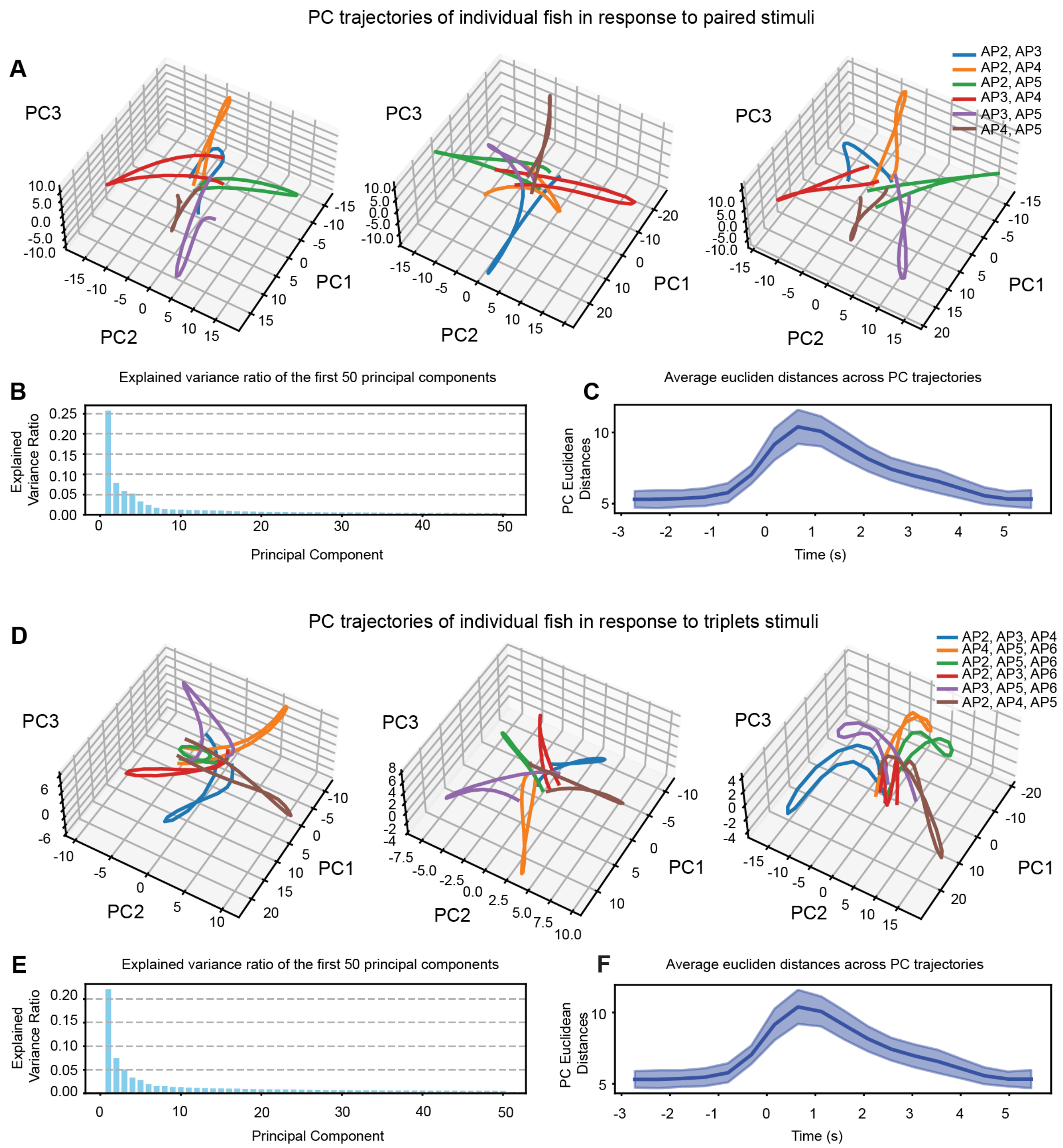


**Figure S10: PCA mean trajectories from individual samples show robust pattern separation. A.** Three examples of the PCA trajectories for three individual larvae exposed to optogenetic stimulation of pairs of neuromasts depict how the population neuronal activity separates when each set of paired stimuli is elicited. **B.** A histogram presents the explained variance ratio of the first 50 principal components from the PCA of the activity matrix of neurons responsive to paired stimulation of neuromasts. **C.** The average Euclidean distance across all PCA trajectories for paired stimuli shows that the maximum distance between the trajectories occurs during optogenetic stimulation. **D.** Like in A., the plot shows three examples of PCA trajectories for three individual larvae exposed to optogenetic stimulation of neuromast triplets, which also demonstrate separated trajectories between stimuli. **E.** A histogram presents the explained variance ratio of the first 50 principal components from the PCA of the activity matrix of neurons responsive to triplet stimulation of neuromasts. **F.** The average Euclidean distance across all PCA trajectories for triad stimuli shows that the maximum distance between the trajectories occurs during optogenetic stimulation.

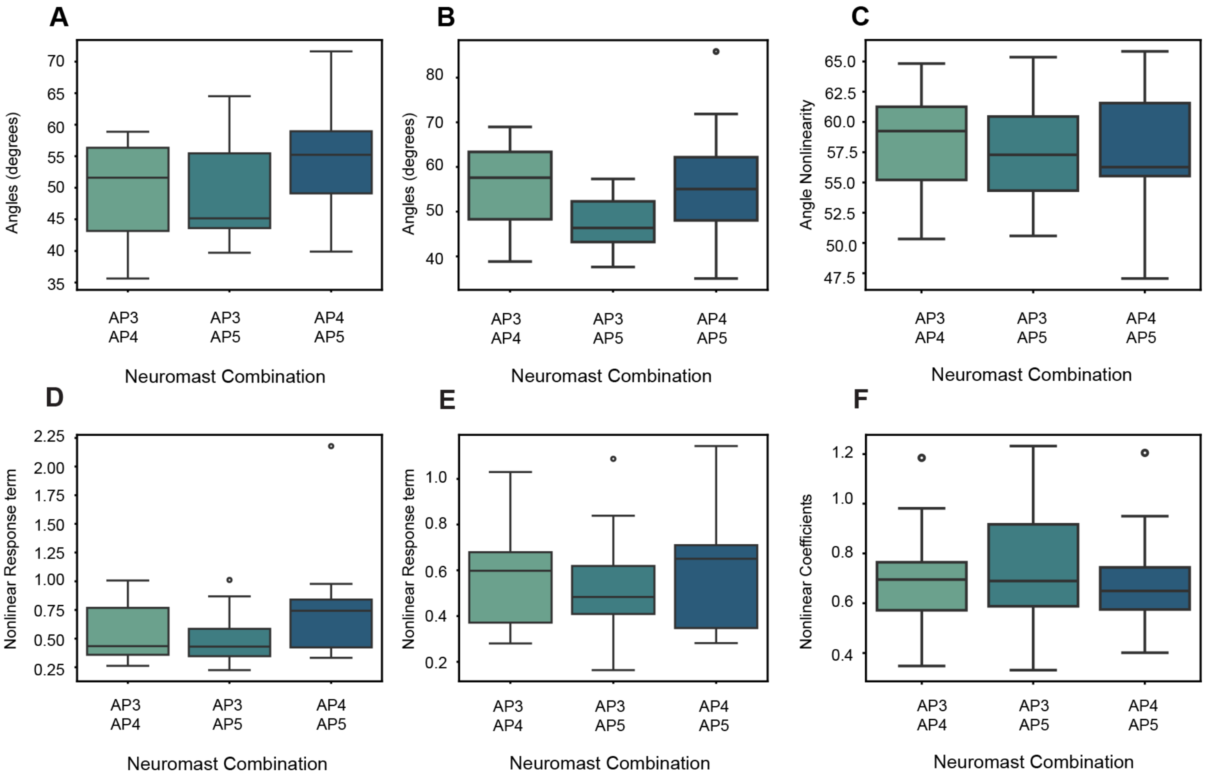


**Figure S11: Null model controls to test the validity of our method to measure the angle of nonlinearity. A and B.** The values of the angle of nonlinearity across pairs of neuromasts, when calculated on non-correspondent pairs, show a higher angle towards orthogonality compared to the angle calculated for correct pairs; however, they do not reach full orthogonality in the *R(A,B)* plane due to residual redundant activations. **C.** The values of the angle of nonlinearity across pairs of neuromasts, when the population activity of neuromast-responsive neurons is scrambled, also indicate a higher angle towards orthogonality. This suggests that the nonlinearities observed are not attributable to the random structure of the neuronal activity being analyzed. **D, E, F.** Three boxplots show the values of the $R_{NL}(A,B)$ residual for the null model control examples in A, B, and C, respectively.

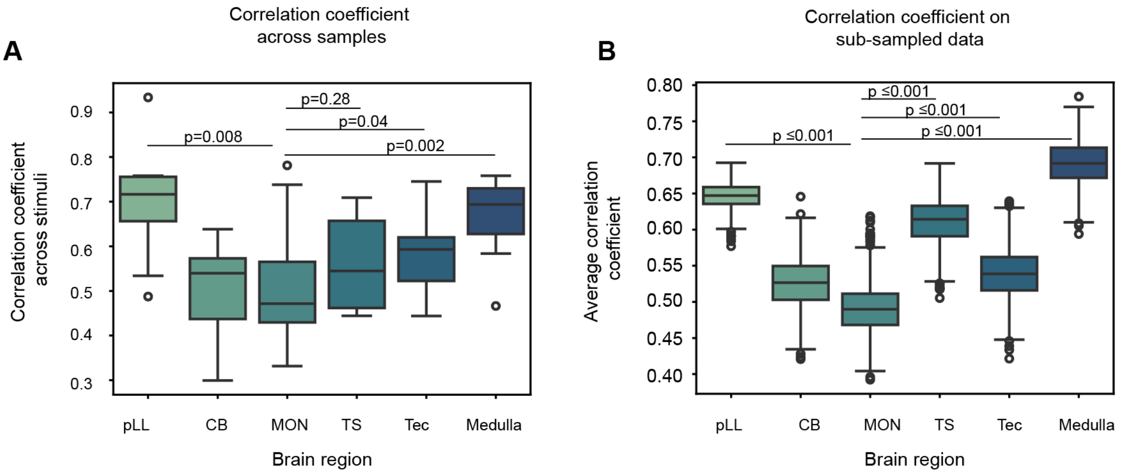


**Figure S12. Average correlation coefficients across samples and when subsampled to address bias in the number of neurons. A.** A graph showing the average correlation coefficients across fish samples for each of six anatomical areas in the zebrafish larva's brain. **B.** Sub-sampled average correlation coefficients across 1000 iterations for each of the six brain regions of the zebrafish larvae (n= 200 neurons) confirm that MON activity during stimulation is more decorrelated than upstream and downstream regions.

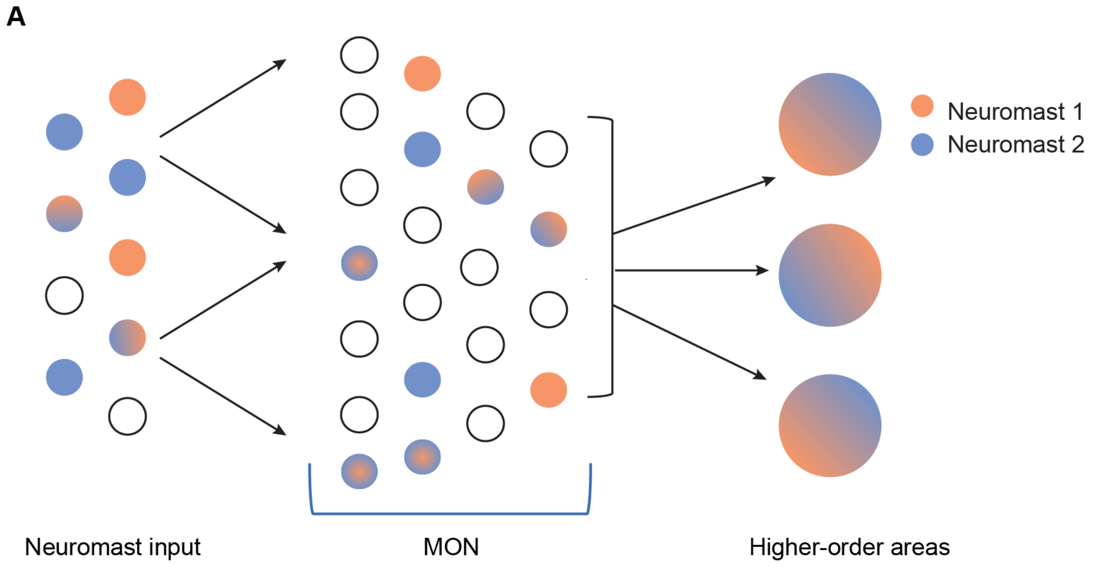


**Figure S13: A working model of the role of higher-order areas in the brain of the 6 dpf zebrafish larva for the discrimination of neuromast input. A.** A schematic that illustrates our working theory that the MON acts as an expanded layer responsible for separating complex neuromast input before distributing pertinent information about it to higher order areas within the zebrafish brain, which generalize set input. In the MON, small and sparse ensembles of neurons enable discrimination among different neuromast input patterns.
